# FLASH Radiotherapy is faster than a heartbeat: A compartmental model to illustrate the interplay between tissue oxygen perfusion and ultra-high dose rate effects

**DOI:** 10.64898/2026.03.12.711443

**Authors:** P. Ballesteros-Zebadua, J. Jansen, V. Grilj, J. Franco-Perez, M-C. Vozenin, R. Abolfath

## Abstract

Ultra-high-dose-rate therapy enhances the protection of normal tissues and reduces side effects while effectively controlling tumors. This biological phenomenon is called the FLASH effect, and when observed, therapy is called FLASH Radiotherapy (FLASH-RT). Various hypotheses have been proposed to explain how ultra-high dose rates achieve these effects under different conditions, with the impact of tissue oxygen perfusion still needing further investigation. FLASH-RT involves brief exposure to radiation, which results in fewer heartbeats occurring during the irradiation period, which could lead to reduced tissue oxygen perfusion occurring during the treatment timeframe. Therefore, we developed a compartmental model to simulate oxygen transfer and its interaction with radiation. The proposed model consists of three compartments: 1) the heart and arteries; 2) the irradiated brain’s blood vessels and capillaries; and 3) the irradiated brain tissue. We employed a system of differential equations, incorporating experimental data from in vivo oxygen measurements using the Oxyphor probe in the brain, to fit the model parameters to the experimental results. This model shows how dose rate and oxygen perfusion could influence chemical processes such as lipid peroxidation, potentially leading to differential biological effects. Our analysis of lipid peroxidation as a function of dose rate revealed a sigmoidal dose-rate-response curve that correlates well with several published biological response datasets. Our results indicate that the differential chemical effects of FLASH-RT compared with conventional dose rates may depend on factors such as oxygen perfusion, consumption, and tissue oxygen tension. This suggests that the temporal dynamics of oxygen could play a crucial role in enhancing the therapeutic window for FLASH-RT treatments. Furthermore, it suggests that the magnitude of some observed FLASH effects may vary across tissues or tumors and across experimental models, given differential oxygen dynamics.

## 1. Introduction

Radiotherapy is one of the main treatments for cancer. However, the use of ionizing radiation to treat tumors is still limited by the adverse effects that radiation dose can induce in normal tissues. In the last decade, dose delivery at ultra-high dose rates has been demonstrated as a groundbreaking technology, capable of producing the FLASH effect, which protects normal tissue from side effects while maintaining tumor control. Interestingly, the exact mechanism of how ultra-high dose rate achieves these differential biological benefits is currently under investigation. (Abolfath et al 2022, 2025, 2020, 2023, Vozenin et al 2026, Limoli and Vozenin 2023)

Oxygen is a well-known radiosensitizer, whereas hypoxic tumors are recognized to be radioresistant (Thoday & Read, 1947). We know now that the mechanisms of oxygen-associated radiosensitisation are intrinsically linked to the radiochemical reactions induced by radiation in the presence of oxygen, and the formation of peroxyl radicals (ROO·) (von Sonntag 1987). These peroxyl radicals can initiate reactions such as peroxidation, damaging membrane lipids and other biomolecules (Phaniendra *et al* 2014).

Interestingly, the role of oxygen tension in the response to ultra-high dose rates has been reported since the first in vitro experiments (Town 1967). In water, FLASH radiotherapy dose rates (FLASH-RT) produce fewer reactive oxygen species (ROS) than at conventional radiotherapy dose rates (CONV-RT) (Montay-Gruel *et al* 2019a, Kacem *et al* 2022, 2025, Abolfath *et al* 2025, 2020). In liposomes and micelles, ultra-high dose rates induce less lipoperoxidation than conventional dose rates (Froidevaux *et al* 2023). However, reducing oxygen tension reduces the difference between ultra-high and conventional dose rates as the reaction of peroxidation is strictly related to the presence of oxygen required to perpetuate the cascade of reactions (Froidevaux et al T2023). In vivo, FLASH-RT dose rates (FLASH-RT) have also been shown to protect normal tissue in physiological conditions (after breathing ambient 21% O_2_). In the brain, FLASH-induced neuroprotection afforded by electron and proton beams is lost when animals are supplemented with carbogen and oxygen (Montay-Gruel *et al* 2019b, Iturri *et al* 2023). Globally, these studies support the idea that the complex interaction between radiation and oxygen is a crucial contributor to the FLASH effect.

Several mathematical models that address variations in oxygen and dose rates have been proposed (Petersson *et al* 2020, Scifoni *et al* 2025, Poulsen *et al* 2024). However, to our knowledge, our model is the first compartmental model to merge mechanistic representations of temporal oxygen perfusion with phenomenological parameterizations of tissue oxygenation, grounded in experimental data.

The duration of irradiation is the most obvious parameter that distinguishes ultra-high dose rates (1 microsecond to 10 milliseconds) from conventional dose rates (2-4 minutes). We hypothesize that the reduced oxygen perfusion of tissues that occurs during FLASH-RT irradiation could trigger the FLASH effect. To explore this hypothesis, we developed a mathematical compartmental model to describe oxygen perfusion in irradiated tissues. We used measurements of oxygen tension performed in the brains of mice during irradiation to set the model’s parameters. We also assessed how the variation in oxygen tension during brain irradiation affects lipid peroxidation in relation to the dose rate. The model shows that tissues can experience reduced oxygen reperfusion during FLASH-RT dose rates, which could influence other processes, such as lipid peroxidation, potentially resulting in lower peroxidation rates for FLASH-RT compared to conventional dose rates. We also showed how tissue- or tumor-specific parameters that can vary across experimental models, such as oxygen consumption, oxygen perfusion, and oxygen tension, can influence the observed dose-rate dependence.

## 2. Methods and Materials

### 2.1 Compartmental Model of Oxygen Perfusion in the Brain

Oxygen perfusion refers to the delivery of oxygen to the tissues through the bloodstream. We propose a compartmental model to illustrate oxygen perfusion, using the brain as an example. The model considers three main compartments (Fig 1): 1. The heart and vessels (arteries and veins); 2. The smaller vessels (arterioles and venules) and capillaries inside the irradiated brain; 3. The irradiated brain tissue.

**Figure 1.**
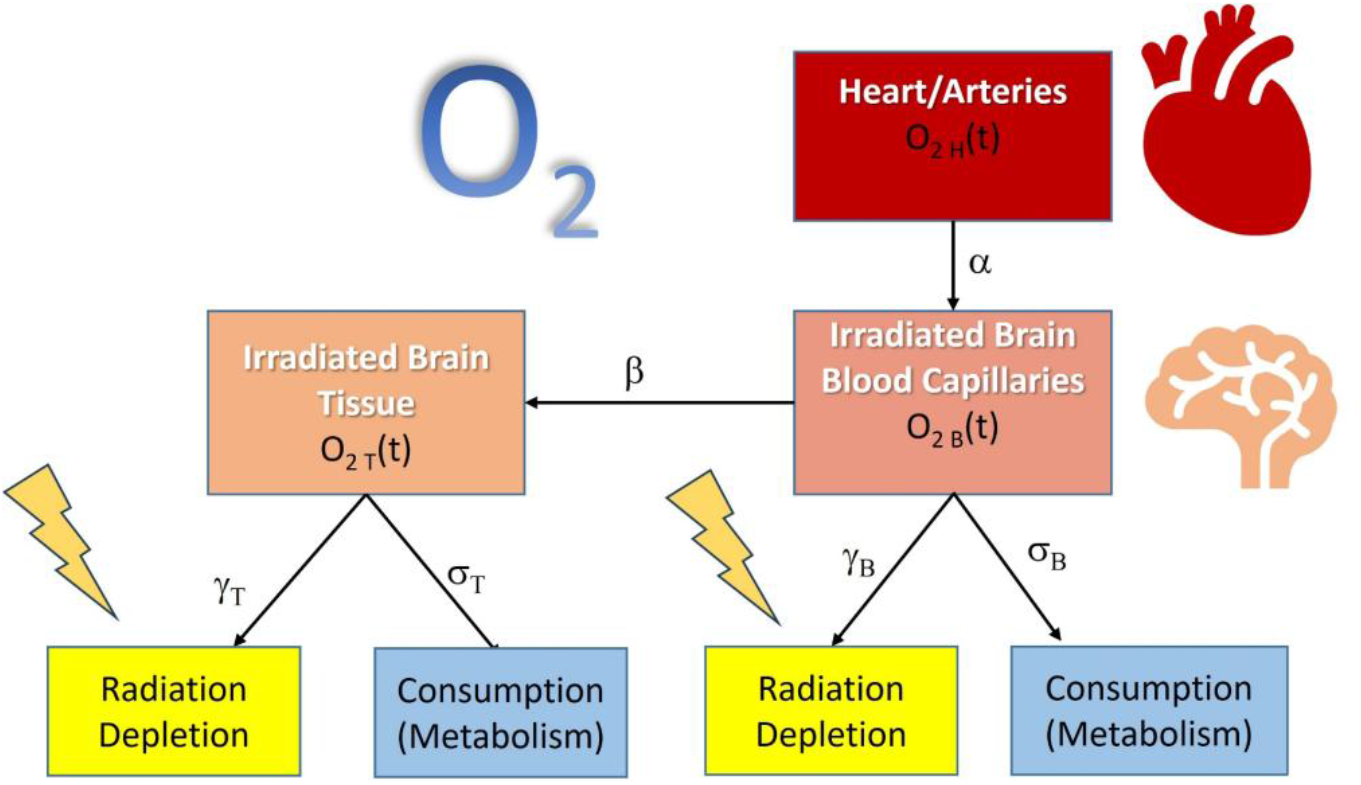
Compartmental Model for the oxygen perfusion in the brain tissues

O_2H_(t) refers to the oxygen being pumped from the heart and arteries with a sinusoidal heartbeat (at a rate α) (Eq. 11), O_2B_(t) refers to the oxygen in the capillaries inside the irradiated brain (arriving at the same rate α) (Eq. 7), and O_2T_(t) refers to the oxygen being perfused inside the irradiated brain tissue (at a rate β) (Eq. 8). Oxygen in the brain is physiologically consumed by metabolism (rate σ_T_) and replenished (rate β) constantly to maintain a quasi-equilibrium state. When the brain is irradiated, a small fraction of the oxygen in both brain compartments is known to be initially fast depleted (with the rates γ_B_=γ_T_=γ) due to the radiochemical generation of free electrons that convert charged neutral oxygen to negatively charged oxygen (e.g. · 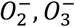 (Labarbe *et al* 2020), owing to direct ionization of biomolecules and/or water molecules in cells..

The initial depletion rate in tissue γ_T_ in the present model depends on the dose rate and was determined by Γ fitting data obtained from oxygen depletion measurements in cell buffer (in press). Then this function is multiplied by a square pulse function P(t) to simulate the radiation beam pulses, and by a function of the existing tissue oxygen tension, *f*_*T*_(*O*_2*T*,0_), that could be considered unity where there is no dependency on the initial pO_2_. The beam pulse structure P(t) was constructed by the superposition of unit step functions. The fitting used for radiation-induced oxygen depletion, γ, as a function of the average dose rate 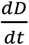 was the following:

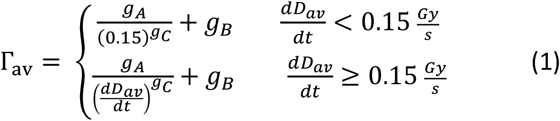

Note that at the limit of 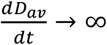 (ultra-high dose rate), Γ_*av*_ → *g*_*B*_. Whereas at the limit of ultra-low dose-rate, 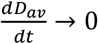, and constant dose, *D*, it saturates to a constant value. In the limit of both negligible dose and dose rate, we expect Γ_*av*_ → 0. In our experimental and theoretical studies, we considered a constant dose (e.g., *D* = 10 Gy). Because below the dose rate cutoff, 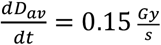, and lack of experimental data for oxygen depletion, and analytical continuation extension of the physical quantities to ultra-low dose rates’s we extrapolated our numerical calculation, assuming *O*_2*T*_ approaches a flat value of 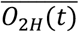 at low dose rates for a given dose value.

A fundamental difference in our notation between the average dose-rate, 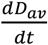, and the instantaneous dose-rate, 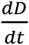 must be realized through time integration of the rate equations. 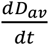 is a parameter, i.e., a constant mean-value, and can be simply separated from the time-integration operator, whereas the time integration over 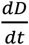 yields the deposited dose within the integrated time interval, hence,

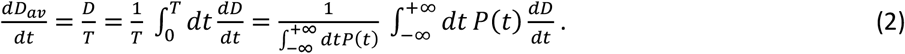

In Eq.(2), *P*(*t*) is the temporal dose rate distribution function. It is important to note that biological endpoints can vary across several orders of magnitude in terms of time scales. Higher than the ultra-high dose rate structures respond to the average dose rate, while the femtosecond dynamics of radiochemical species are sensitive to the instantaneous dose rate.. However, the biological responses can be sensitive to the instantaneous dose rates at ultra-low dose rates where the dose rate time scale extends over biological processes.

The beam pulse structure is defined as P(t). One example of pulse structure using a step function is shown in Fig. 2. Here s is the spacing between pulses, N is the number of pulses, and w is the pulse width (Fig. 2).

**Figure 2.**
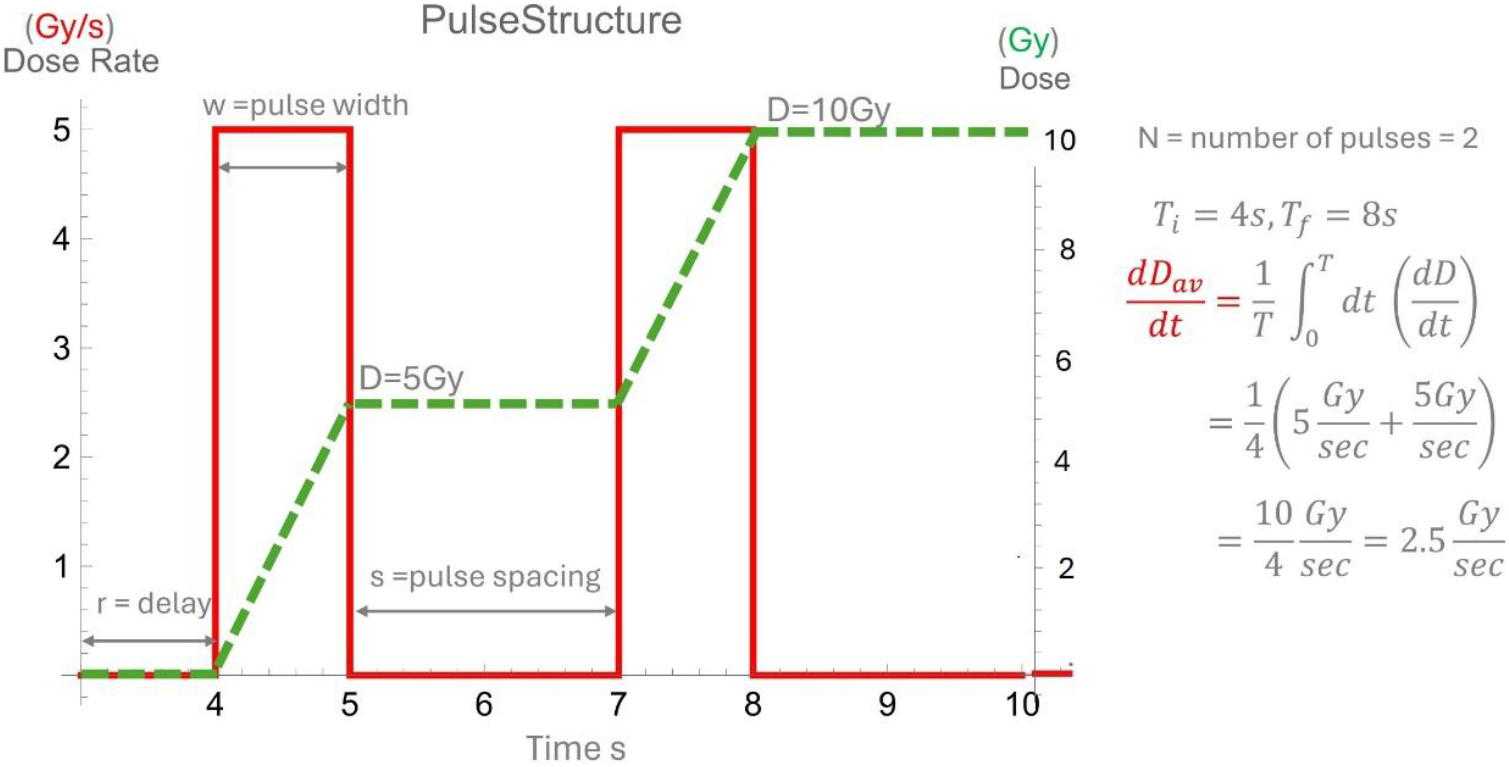
Example of a pulse structure definition P(t) using unit step functions

The initially produced • *OH* by radiation is then considered to interact with the available oxygen, O_2T_(t), and lipids, L(t) in the tissue to initiate radiation-induced lipid peroxidation (LOOH), as outlined in the following chemical reactions.

### 2.2 Chemical Reactions

We consider the following chemical reactions after irradiation (Grilj *et al* 2024b).

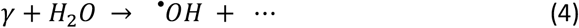

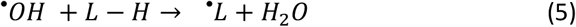

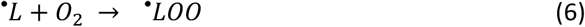

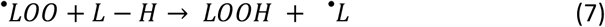

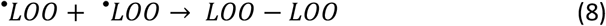

The reaction rate of Eq.7 is significantly slower than that of the other three. Thus, we isolate it and treat it differently in our numerical analysis of the reaction rates, which is presented below.

### 2.2 Oxygen Rate Equations

We used the following system of coupled differential equations to model the oxygen transfer between compartments and their constraints:

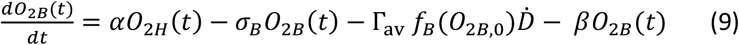

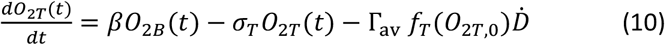

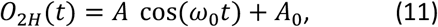

*where ω* = 2*πv, v* = heart rate, and 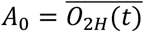

Note that the mean value of 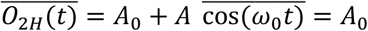, where 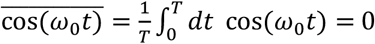 whereas 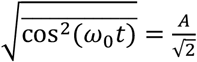 since

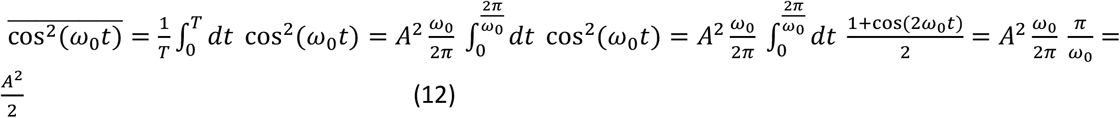

### 2.3 Lipid Peroxidation Rate Equations

In addition to the above equations, we used the following system of coupled differential equations to model the generation of • *OH* and lipid peroxidation, LOOH, and its constraints:

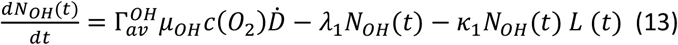

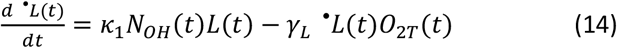

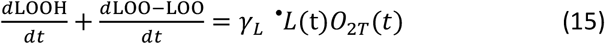

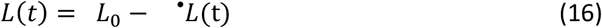

In our numerical solutions of these rate equations, we consider a square pulsed dose rate, given by *P* = *θ*(*t* − *t*_1_)*θ*(*t*_2_ − *t*), thus *T* = *t*_2_ − *t*_1_. As specified above, the instantaneous dose rate and the mean dose rate are denoted by 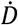, and 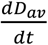.

*μ*_*OH*_ is the mean ^•^*OH* G-value, expressed in M/Gy (Parameter values in Supplementary Data Table 1). 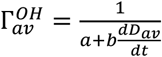 is a prefactor similar to prefactor in oxygen dose rate dependencies, Γ_*av*_, extracted from previous analysis(Abolfath *et al* 2025, 2020), and *c*(*O*_2_) describes the oxygen dependence of ^•^*OH* G-value due to the competition channel with oxygen and recombining •*OH* with free electrons and the formation of *OH*^−^ or any anion species of oxygen (*e. g*., 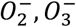). *κ*_1_ the rate at which ^•^*OH* is consumed in the reaction in Eq. 13, and *λ*_1_ is the rate at which ^•^*OH* is consumed in other reactions (Eq 13). *γ*_*L*_ is the rate of ^•^*LOO* generation in the presence of oxygen in the reaction in Eq. 30. In Eq.(14), ^•^*L*(*t*) is coupled with *O*_2*T*_ (*t*). Numerically, we calculate the average dose rate, 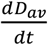 outside of the rate equation loop, as sketched in Fig. 2.

We used a heart rate *ν* of 400 beat per minute (bpm) for a mild anesthetized mouse (Ewald et al., 2011). To obtain the values of *α*, *β, σ*_*T*_ and *σ*_*B*_, we fitted experimental data on oxygen measurements performed in vivo in the mouse brain during FLASH-RT (Grilj *et al* 2024a). After fitting the experimental data, we solved the equations using a personalized numerical code in Fortran to obtain the oxygen values for the brain, and then calculated the amount of lipid peroxidation (LOOH) for different dose rates.

To explain the time integration of *N*_*OH*_(*t*) in Eq. (13), first let us consider the following simplified rate equation

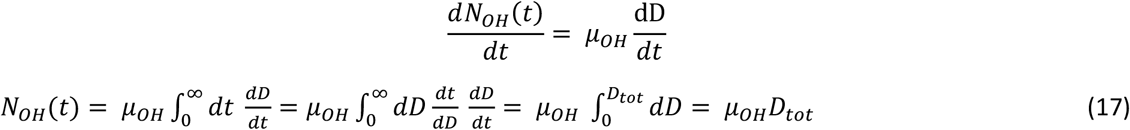

As fitted in equation 1, we know that initial oxygen depletion depends on the dose rate. Depletion is known to be related to the oxygen consumed in radiochemical production, so the higher the free radicals produced by radiolysis, the higher the oxygen consumed. We assume then that free radicals’ production will be dose rate dependent accordingly. This has also been confirmed in the measurements done by Kusumoto et al (Kusumoto *et al* 2024). Therefore, we defined the effective yield 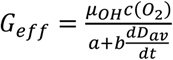 as a function of the averaged dose rate that does not enter into the time integration. 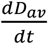 is a constant that only re-normalizes the · *OH* production yield *μ*_*OH*_. Hence

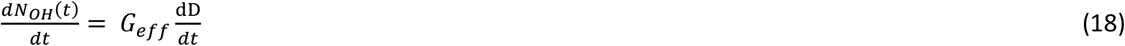

thus

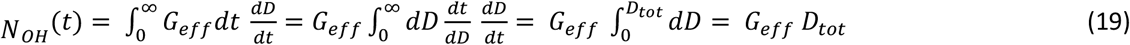

Finally, from equation 15

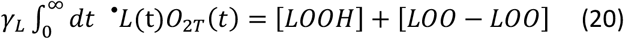

as LOOH and LOO-LOO are considered the end-point compounds.

### 2.4 Analytical and numerical derivation of the dependence of O_2T_ as a function of the initial condition of O_2T_ (t=0)

In the above equations, the initial condition for *O*_2*T*_(*t* = 0) = *A*_0_ coincide with *O*_2*B*_(*t* = 0) = *A*_0_. We, however, know that oxygen tension in the brain tissue is lower than in the blood so we can consider slightly different initial conditions for *O*_2*T*_(*t* = 0) = *A*′_0_ = *ηA*_0_ with 0 < *η* ≤ 1 to investigate the dependence of the solutions as a function of *O*_2*T*_(*t* = 0). Using these initial conditions should lead to the same conditions for the steady state solutions, *O*_2*T*_(*t* = ∞) and *O*_2*B*_(*t* = ∞). In our numerical approach, to avoid numerical instabilities we started from the initial conditions *O*_2*B*_(*t* = 0) = *O*_2*T*_ (*t* = 0) = 0 that leads to the steady state solutions 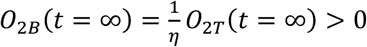. Once we have reached the steady state conditions numerically, we apply the radiation.

The rate equations comprise two dynamical degrees of freedom, corresponding to fast and slow time variations, which represent the oscillatory oxygen pumping from the heart and the oxygen tension in the tissue. Like any other dynamical system with fast and slow time variations, we can integrate over the fast oscillations to obtain an effective dynamics for a slow varying degree of freedom. This is equivalent to the “adiabatic” approximation for the envelope function with time averaged over fast varying degrees of freedom applied in various dynamical systems in physics, such as phase and group velocities of the superposition of two waves with slightly different wavelengths. Note that time averaging over the heartbeat alone, is not equivalent of the time averaging on O_2B_ or O_2T_. As there are fast and slow time scales, we only keep track of the slow evolution of O2B and O2T, through their rate equations by integrating over fast oscillations of O2H, which is integration over small and fast modulations of Heat beats superimposed to large and slow time varying of O_2B_ and O_2T_. We loosely call this operation a mapping to an “envelop” function to differentiate the mathematical operation from time averaging, which is equivalent to smoothening the time evolution of *O*_2*B*_ and *O*_2*T*_ and filtering out small oscillatory modulations due to the heartbeat.

Such dependencies over the envelop functions of *O*_2*B*_and *O*_2*T*_ can be obtained numerically after integrating out the rapid oscillations in *O*_2*B*_ and *O*_2*T*_. For simplicity, we disregard the radiation terms in the rate equations in this section. After performing the time averaging over rapid oscillations (*ω*_0_), in the rate equations (9) and (10), we obtain the following rate equations for the time-averaged functions. Let us first look at the equation for “Blood” where

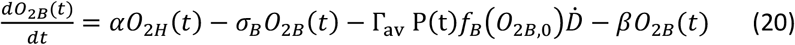

After averaging out the rapid oscillations

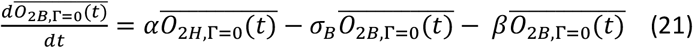

The steady state solution 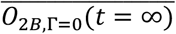 which is the solution of 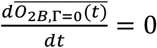 can be found after applying the following 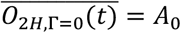. Insertion of this condition in the above rate equation gives

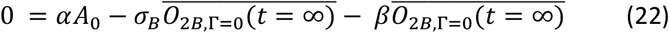

hence

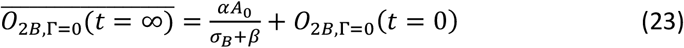

after adding the initial condition, *O*_2*B*,Γ=0_(*t* = 0).

To check, we calculate the time-dependent solution for the envelop function, 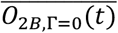 analytically.

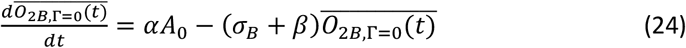

Converting this differential equation to an integral equation

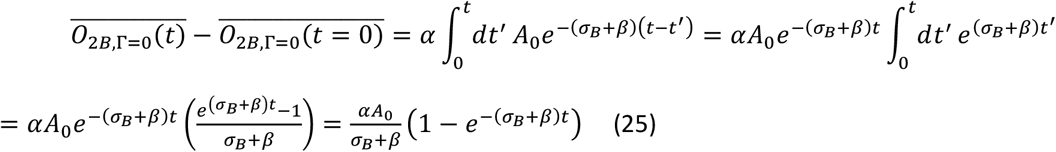

Thus

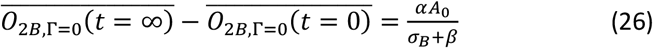

We now turn to the Eq. 10 for tissue:

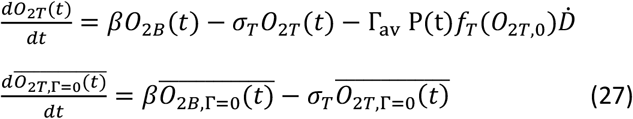

where 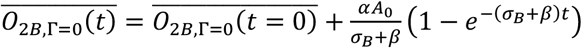. Converting this differential equation to an integral equation:

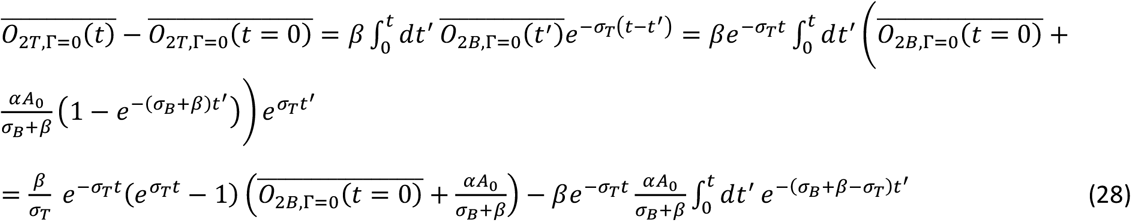

Hence,

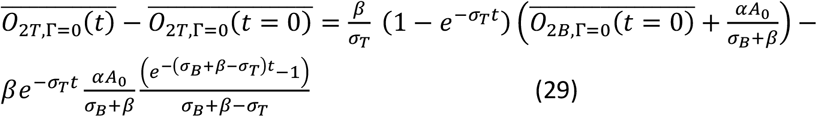

Finally, rewriting,

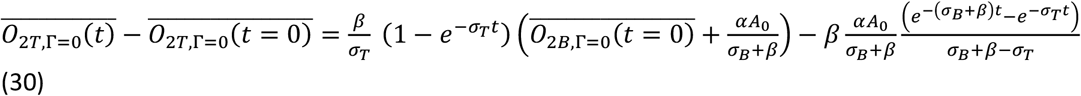

In summary,

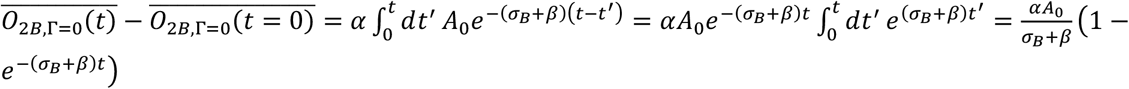

where 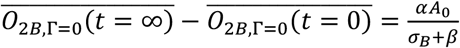, and

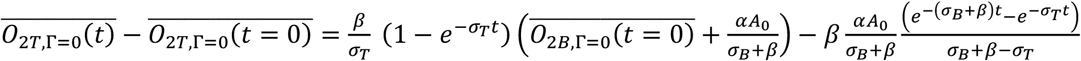

Where 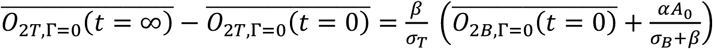.

### 2.5 Solutions of the rate equations in the presence of a field of radiation

We denote the 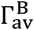 and 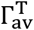 for the radiation effect on oxygen in the blood and tissue. The rate equation for the envelope functions in the presence of continuous radiation can be written as

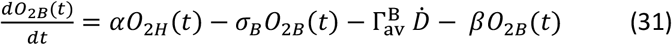

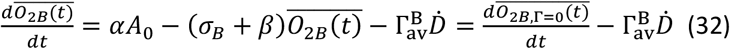

Converting this differential equation to an integral equation

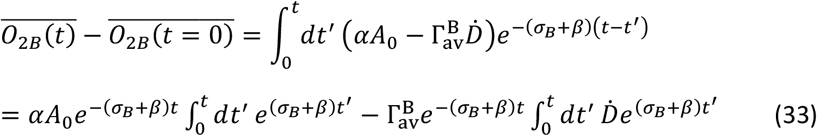

The first term in the right-hand side of Eq. 33 corresponds to zero radiation, 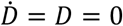. Hence

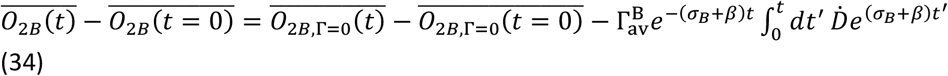

The last term on the right-hand side of Eq. 33 can be calculated using the square pulse approximation. For a square dose rate, 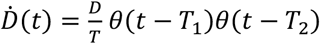 for *T*_1_ ≤ *t* ≤ *T*_2_ where *T* = *T*_2_ – *T*_1_, 

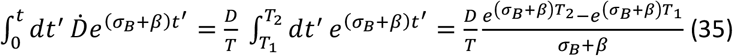

Thus

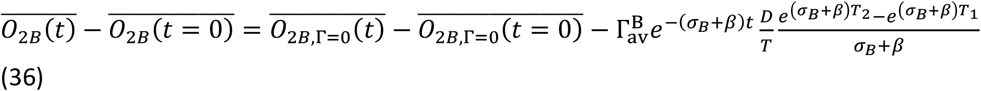

The *θ*(*t* − *T*_1_)*θ*(*t* − *T*_2_) gives a time interval during which the field of radiation was applied. The last term on the left-hand side can be rewritten as

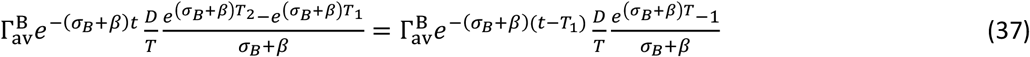

*t* − *T*_1_ is a shift in time, which can be handled appropriately if there were numerical issues due to the large *t* limits. In particular, if we would like to consider radiation in the steady state time domains, then *T*_1_ must be a large number. That causes the numerical solutions to diverge from the exponentials.

Previously we calculated

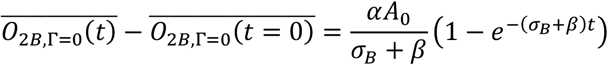

Thus

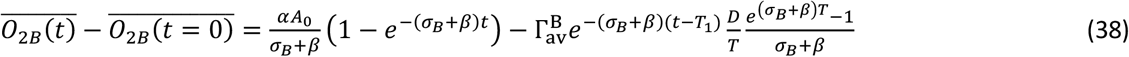

We now turn to the equation for tissue:

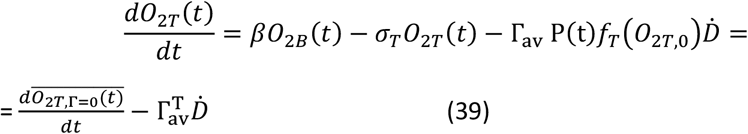

And follow the same rule of thumb:

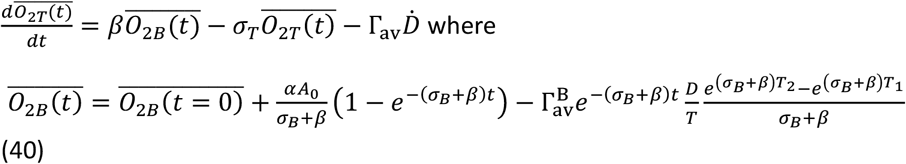

Converting this differential equation to an integral equation:

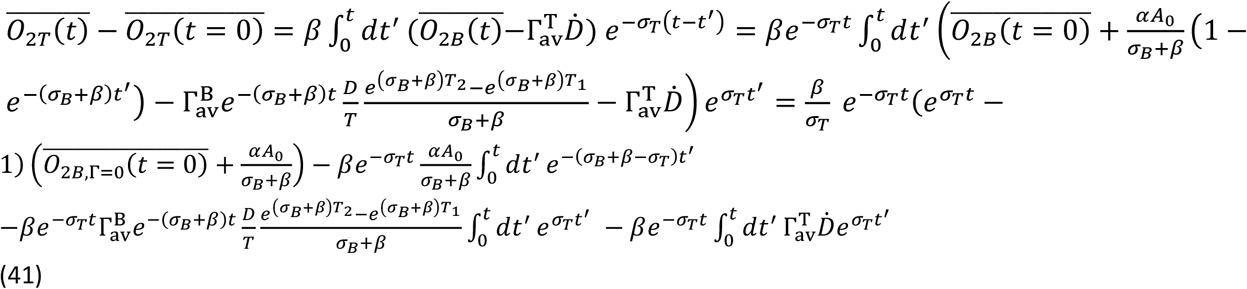

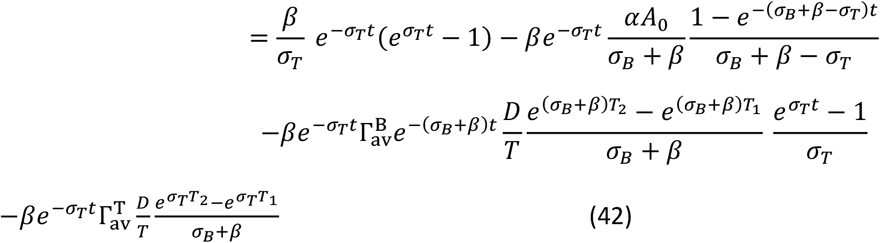

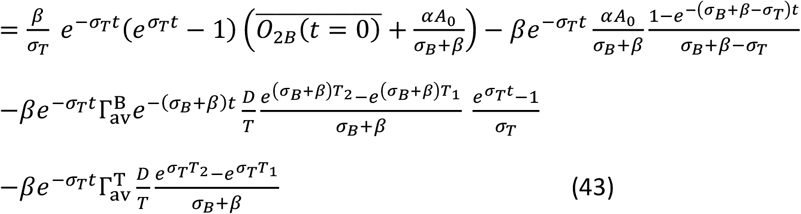

where

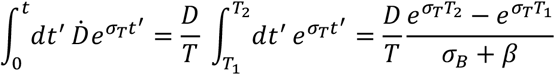

Finally

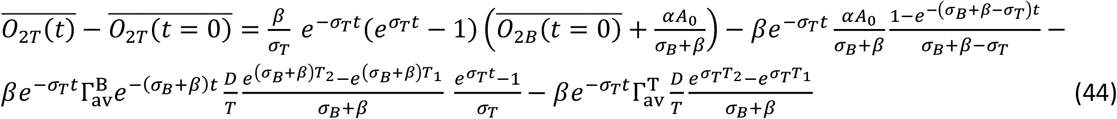

Again, we can rearrange the radiation terms on the right-hand side to avoid the floating-point divergence of the exponential in numerical calculation

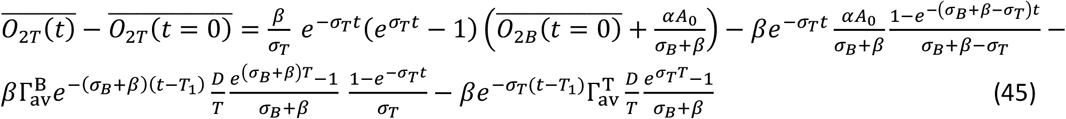

The steady state solution is

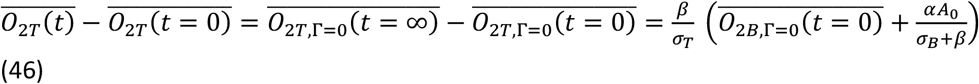

In summary,

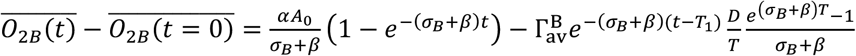

where 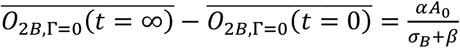, and

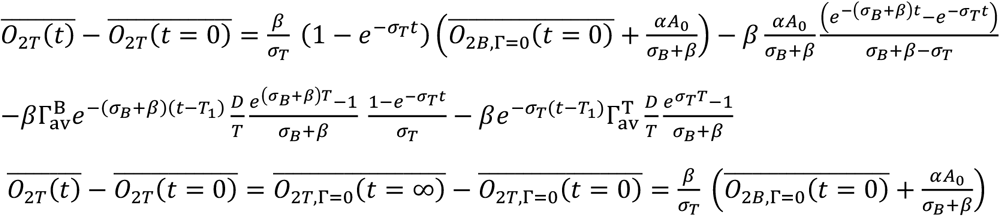

Note that

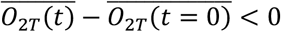

is the oxygen depletion.

### 2.6 Numerical calculations and divergence

Due to the large *T*_1_ we may face the floating divergence in our numerical calculation. To handle that, we run the numerical code using two split solutions at *t* ≤ *T*_1_ and *t* > *T*_1_ and matching the solutions at *t* = *T*_1_

*t* ≤ *T*_1_:

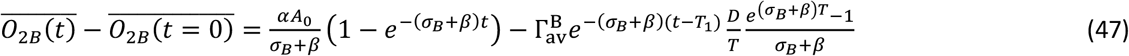

and

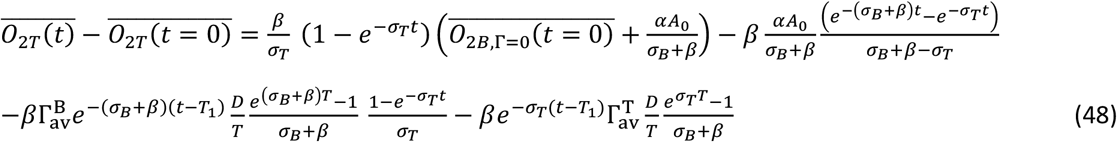

Using *τ* = *t* − *T*_1_, hence *τ* > 0:

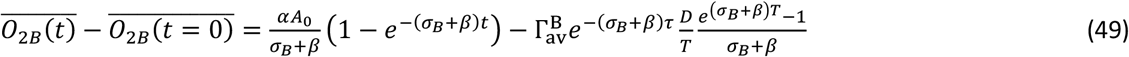

and

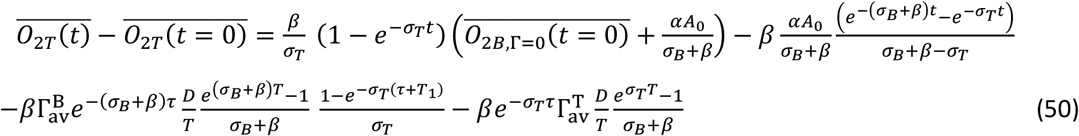

For the beam pulse and ultra-high to conventional dose rate threshold, **w**e start with Eq. 50

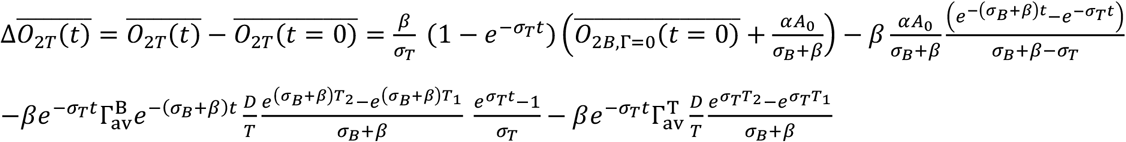

where 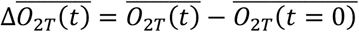. Considering constant dose, *D*, the variable that controls the dose rate is *T* = *T*_2_ − *T*_1_. We first change the variable *T*_2_ = *T* + *T*_1_

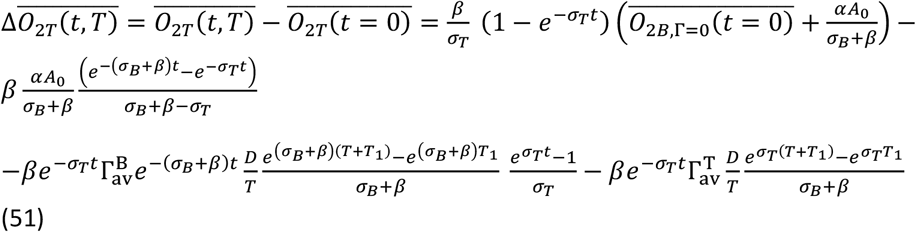

Let us take a derivative with respect to *T*

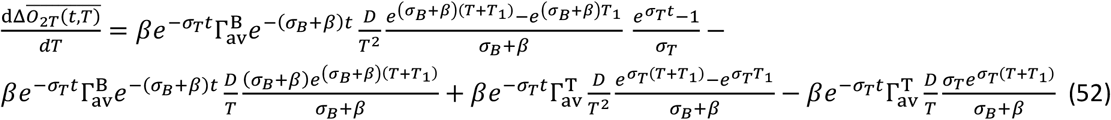

The optimal *T* can be found by assertion of the condition 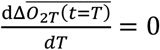 at *T* = *T*^∗^ where T* is the optimal pulse width that, for a continuous beam determines the optimal dose rate obtained as follows

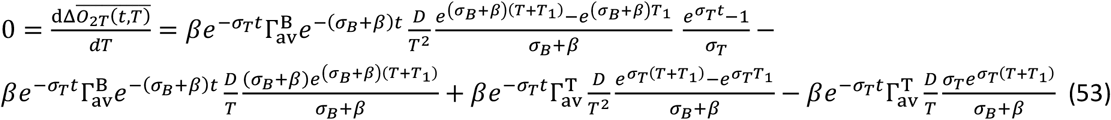

Simplifying this equation

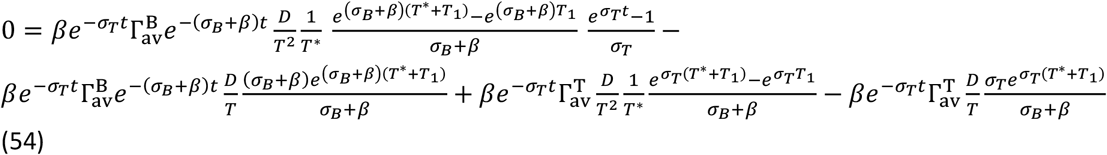

Considering *t* = *T*_1_

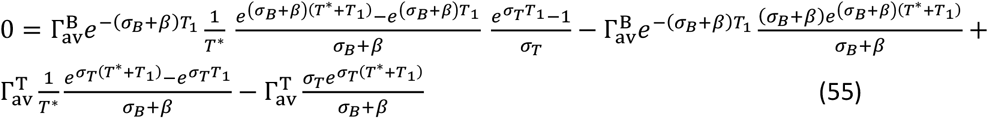

Factoring 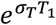

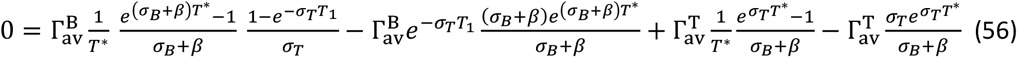

Using *T*_1_ → ∞ we find the solution for *T*^∗^ in the following

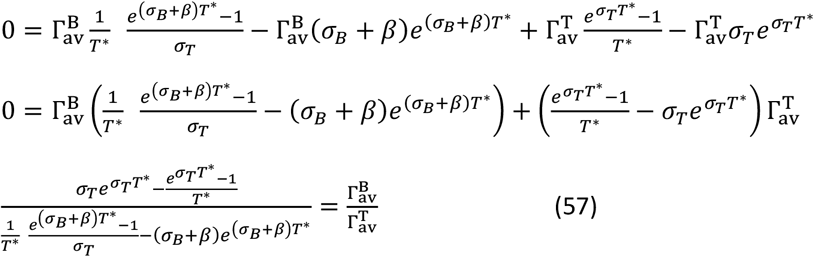

*T*^∗^ can be calculated iteratively from the above equation. This implies that the optimal ultra-high dose rate with the highest differences from ultra-high dose rates from conventional dose rates is the following

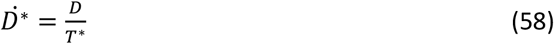

where the irradiation time is faster than the time needed to re-oxygenate.

### 2.7 Numerical Solutions

Recalling

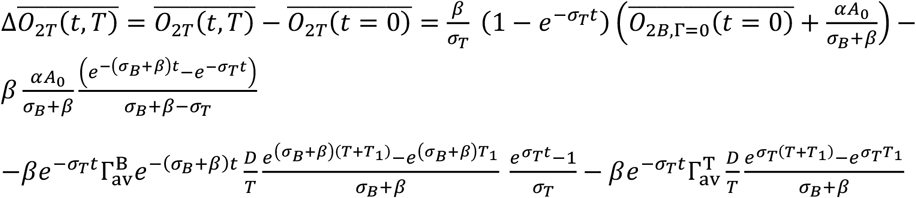

And using *t* = *T*_1_, and 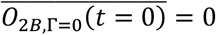

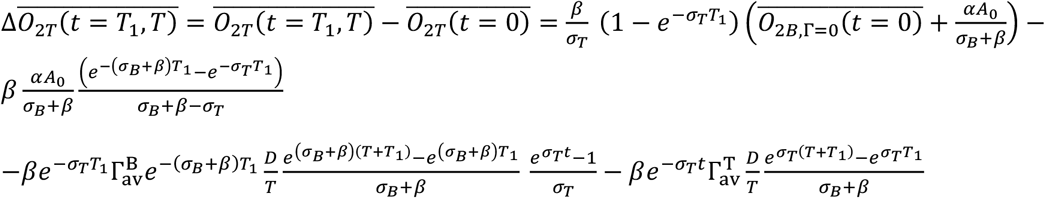

At the steady state time interval, where *T*_1_ = ∞

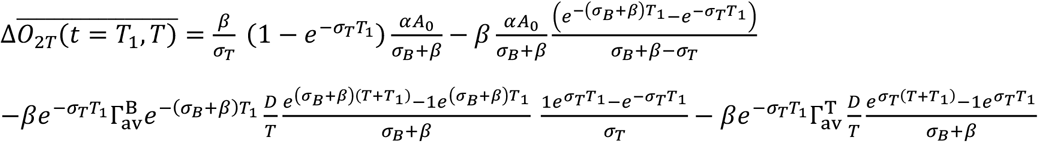

That is

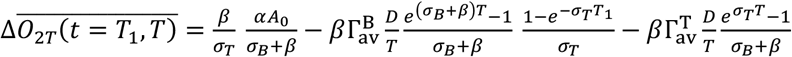

For *T* → 0:

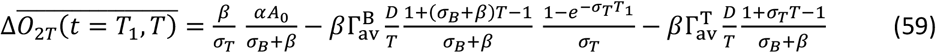

### 2.8 Tissue Oxygen tension and dose rate

To introduce the difference in initial oxygen tension between tissues, let us recall the solutions of the rate equations

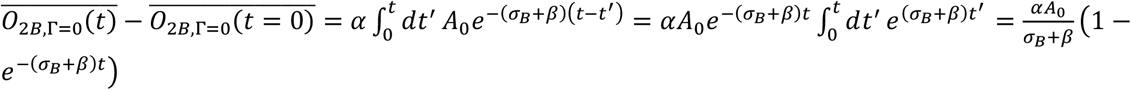

where 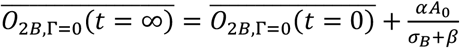, and

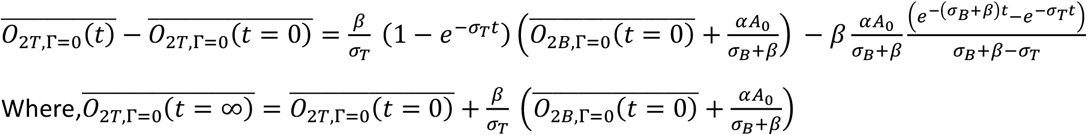

Where, 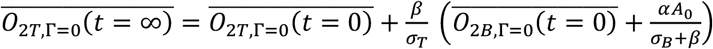

Let us subtract the steady state

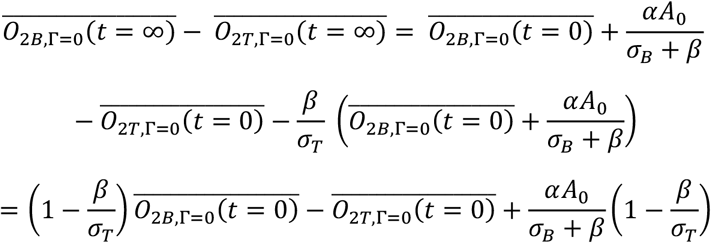

Considering 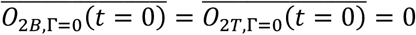 as the initial condition for the differential equations, we obtain

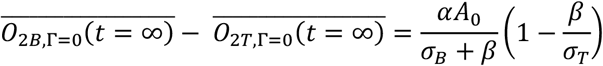

To have 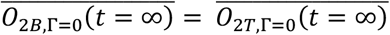, an expected oxygen condition in tissues like the heart, it is sufficient to set 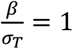. This condition is, however, suitable for tissues with an equilibrium between perfusion and consumption in normal conditions.

Introducing 0 ≤ *η* ≤ 1, for tissues with less oxygen than the arterial level (e.g. PO2 in the arteries in the brain is 100 mmHg, and in the tissue around 40mmHg) we expect 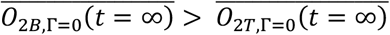, hence we introduce 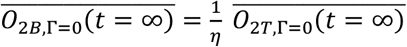. Note that, mathematically, the steady-state solutions of the differential equations coincide for high- and low-oxygen-tension tissues, but not the initial conditions, since we deliberately started from zero, which was suitable for mathematical control of the solutions. Thus

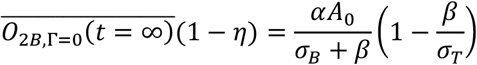

or

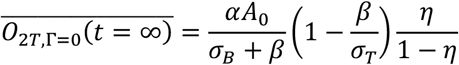

Assuming

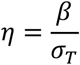

we find

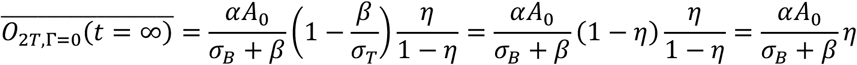

In summary, for 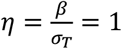, a condition that applies to tissues in physiological equilibrium conditions.

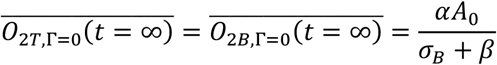

and in a general situation with 0 ≤ *η* < 1, corresponding to tissues with unbalanced oxygen-consumption conditions, like some tumors, we have

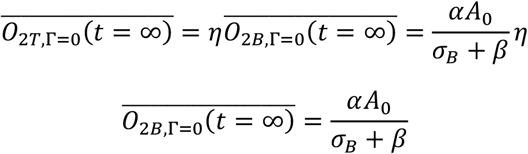

## 3. Results and discussion

### 3.1 Oxygen reperfusion in the brain varies with the dose rate

Initially, the coupled differential equations were numerically solved, and the parameters were optimized to fit a set of measured oxygen tensions in a mouse brain, where the initial brain pO_2_ 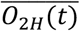 was set at approximately 40 mmHg (38-45 mmHg range) (Grilj *et al* 2024a). By this, the model parameters alpha, beta, and eta were determined based on biological data. For visualization, the model solution was plotted together with the in vivo data. (Fig 3).

**Figure 3.**
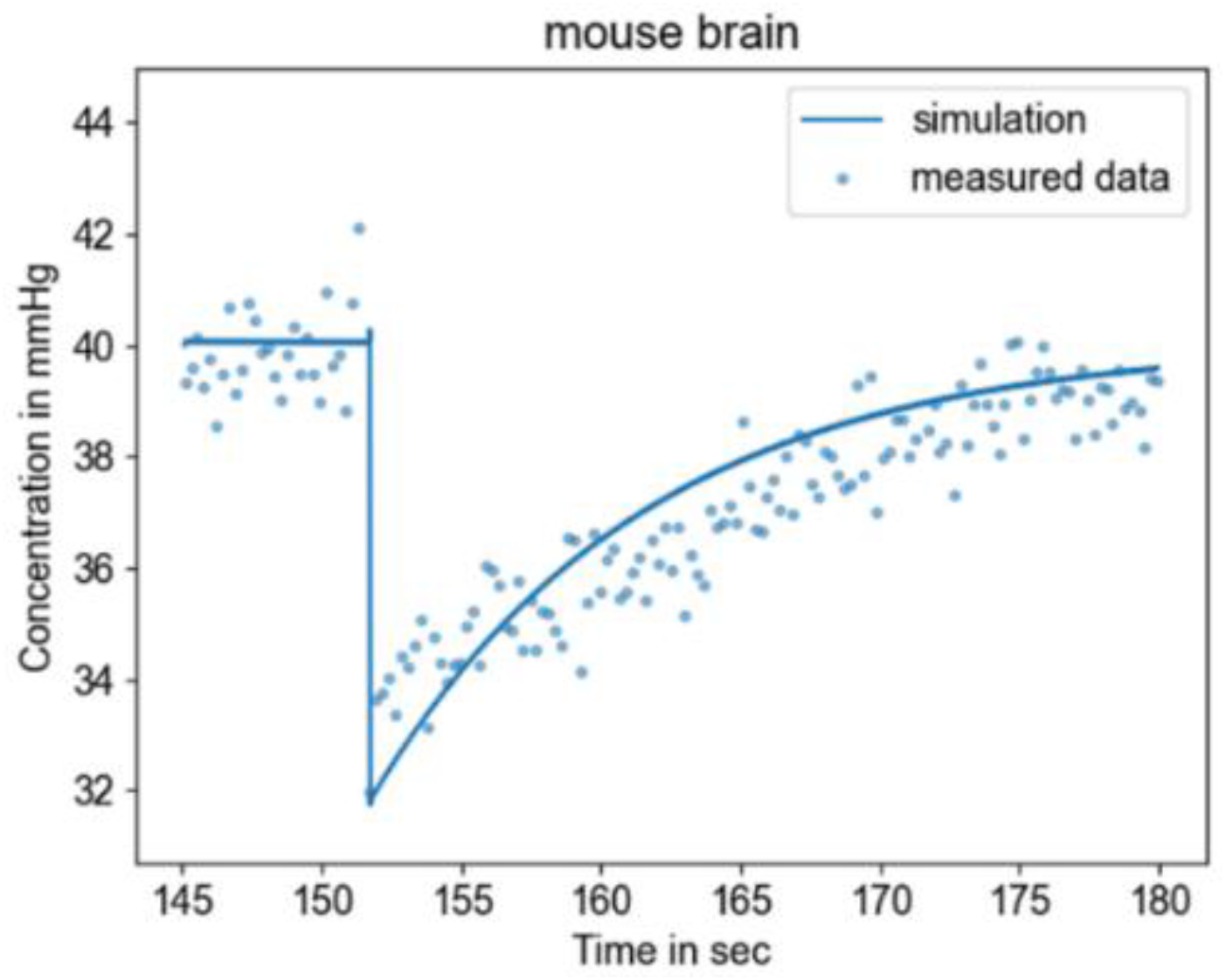
Measured data of pO_2_ (dots) in the mouse brains in vivo and simulated data (line) for one continuous pulse of 10 Gy at 5.5 × 10^6^Gy/s according to the compartmental model differential equations after fitting rate parameters.

The solutions of the rate equations were obtained numerically to simulate the oxygen behavior in the blood and the tissue after irradiation (Fig. 4A) for one pulse of 10 Gy delivered whole brain, we can observe an immediate drop in oxygen levels from radiolytic oxygen depletion (ROD) followed by an oxygen replenishment phase (O_2T_) due to perfusion of oxygen and pumping up by blood vessels in the tissue. It is important to note that decreasing the heart rate (e.g., from mice at 400 bpm to humans at 80 bpm) has a small impact on tissue oxygen, which is dominated by the perfusion coefficient. Fig. 4B shows the calculated formation of radicals (Fig. 4B) where the production yield of radiation-induced ^•^*OH*-radicals *in vivo* can be obtained by d *N*_*OH*_/dt at time zero (irradiation time).

**Figure 4.**
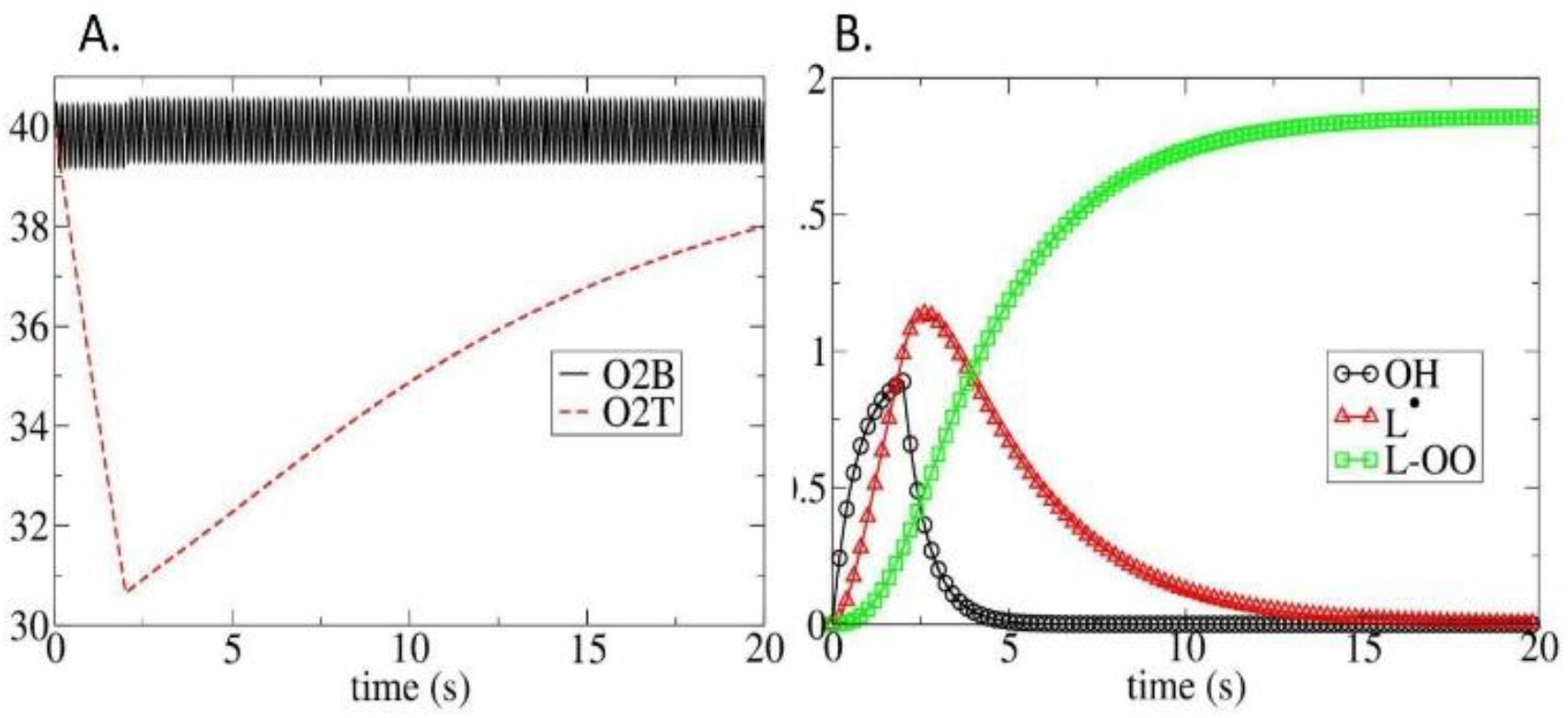
Numerical solutions of the rate equations (Eq. 9-11, 13-16) for one pulse with a dose of 10 Gy in the brain at a dose rate of 5Gy/s, showing A. Oxygen oscillations in blood vessels (O_2B_ - black) are superimposed with a drop in oxygen level in the tissue (O_2T_- red) due to radiation-induced oxygen depletion. B. Time evolution of ^•^*OH*, ^•^*L*, and ^•^*LOO* radicals created after irradiation.

To evaluate how oxygen is reperfused in the tissue as a function of the dose rate, we ran simulations at various dose rates, as illustrated in Fig. 5. The amount of oxygen replenishment occurring during irradiation varies significantly with dose rate and tissue parameters. Here, we see that when we move from conventional dose-rates (yellow) to ultra-high dose rates (red), the drop in oxygen appears steeper and more pronounced for the ultra-high dose rates because less oxygen is perfused during the shorter irradiation time. Therefore, for ultra-high dose rates, the drop is observable, whereas for conventional dose rates, it is less apparent, as oxygen depletion is compensated by reperfusion during irradiation. This is consistent with previous *in vivo* experiments where oxygen depletion was not measurable after irradiation at conventional dose rates due to oxygen reperfusion(Cao *et al* 2021, Grilj *et al* 2024a).

**Figure 5.**
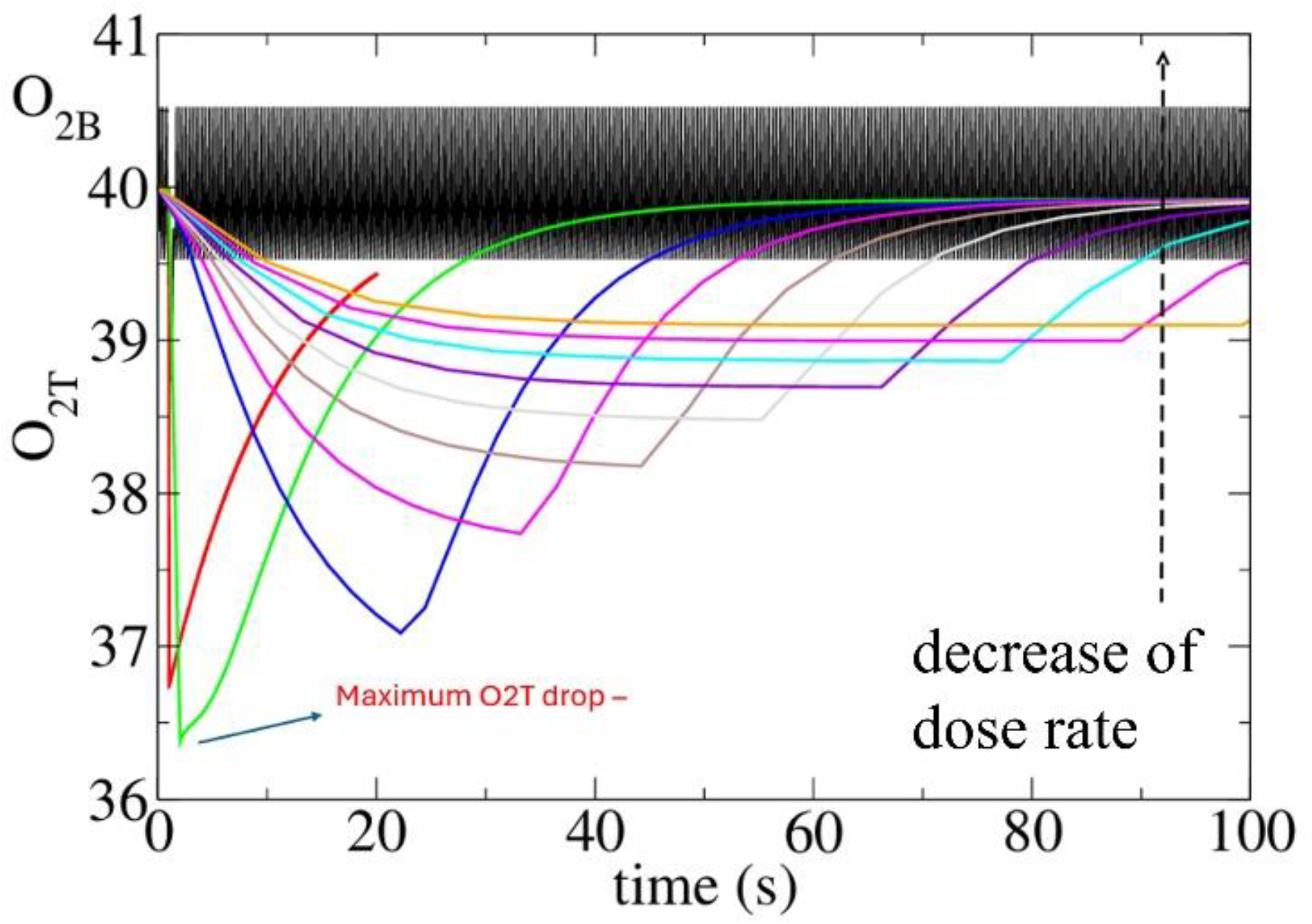
Time evolution of O_2B_ (black) and O_2T_ (colors) in the brain as a function of time and dose rates for a given continuous dose of 10 Gy. The average dose rate ranges from 500 Gy/s to 0.1 Gy/s. The first dose rate (red) is 500 Gy/s, the second dose rate (green) is 10 Gy/s, and the third one is 0.45. 0.30, 0.22, 0.18, 0.15, 0.13, 0.11, 0.1 Gy/s. That is for 10 curves.

In figure 5, interestingly we can find a value where the influence of reperfusion is minimal and the observable depletion is the tissue maximum, as seen in Eq. 58. According to this equation, the dose rate value at which this maximum drop in oxygen will be observable depends on the total dose, but also on perfusion and consumption and depletion parameters defined in the model. Using the parameters obtained from the experimental brain data, the maximum drop observable occurs at a dose rate of approximately 10 Gy/s. According to these first results and for further comparisons we will use FLASH-RT to refer to time averaged ultra-high dose rates > 10Gy/s. Similarly, we will define CONV-RT to further refer to time averaged dose rates < 1 Gy/s.

### 3.2 Initial tissue Oxygen tension and dose rate

To calculate how the initial level of oxygen in the tissue may influence the oxygen dynamics after irradiation in the tissue, we ran simulations at different initial values of oxygen tension, modifying the value of eta arbitrarily to evaluate its influence, as illustrated in Fig. 6. Here, we can clearly observe that before radiation, changing the parameter *η* also modifies the initial tissue oxygen level, but the shape of the observed change in tissue oxygen after irradiation is preserved.

**Figure 6.**
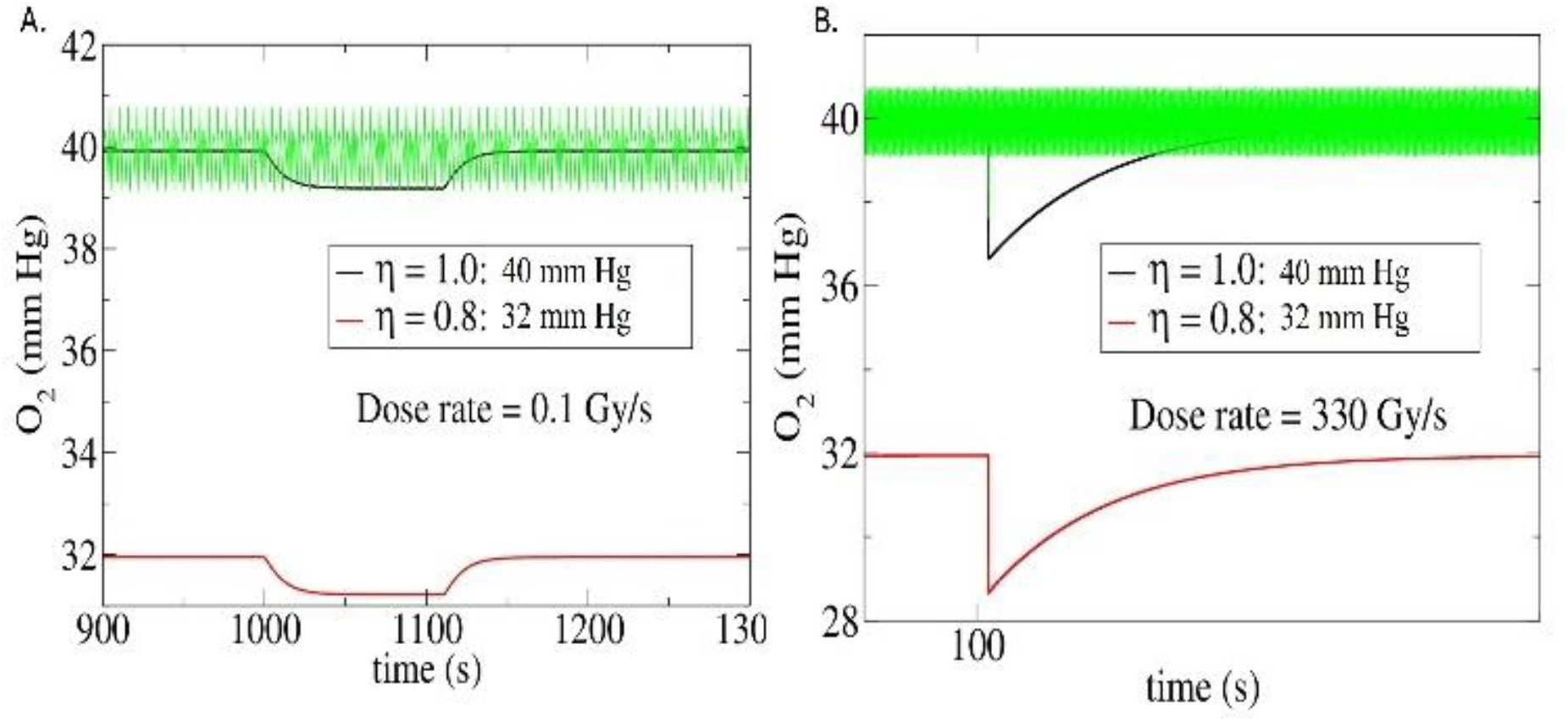
Numerical solutions of the rate equations (Eq. 9-16) for one continuous pulse with a dose of 10 Gy in the brain at two different average dose rates of A. 0.1 Gy/s and B. 330 Gy/s. showing oxygen oscillations in blood vessels (green O_2B_) superimposed on a drop in tissue oxygen level (O_2T_) due to radiation-induced oxygen depletion. Black lines represent an initial oxygen tension of 40 mmHg and a well-balanced eta of one *η* = 1. Red lines represent a lower initial oxygen tension of 32 mm Hg and a less balanced eta *η* = 0.8.

To assess how oxygen is reperfused in the tissue as a function of dose rate, for the same eta values, we ran simulations at two different dose rates, as illustrated in Fig. 6 (CONV-RT 0.1Gy/s and FLASH-RT 330 Gy/s).

In Fig. 6, we can also see that when we move from CONV-RT (A. left) to FLASH-RT (B. right), the drop in oxygen is steeper and more pronounced as less oxygen is perfused during the irradiation time, as seen in Fig. 5. So the dose rate influences the shape of oxygen behavior including reperfusion while eta is more related with the initial oxygen value.

### 3.3 How does lipid peroxidation depend on dose rate?

We evaluated LOOH as a candidate that may be differentially altered in response to dose rate (Eq. 15). The oxygen dependence of the OH-radical rate equation (Eq. 13 and Eq 17) was defined as follows:

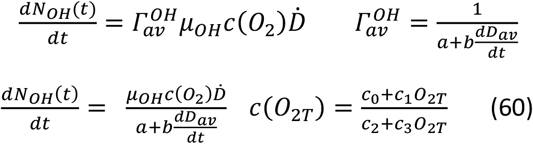

We calculated the area under the curve for LOOH, calculating the integral 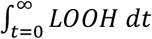 to evaluate the induction of lipid peroxidation by radiation (Fig. 7). When using parameters of oxygen perfusion *β* and consumption *σ*_*T*_ obtained from the mouse brain data, lipid peroxidation is consistently lower at the highest dose rates. Interestingly, lipid peroxidation flattens at average dose rates above 100 Gy/s and higher. Similarly, lipid peroxidation flattens for average dose rates lower than 1 Gy/s. The greatest change of lipid peroxidation with dose rate happens between 1Gy/s and 100Gy/s at intermediate dose rates. The reduced lipid peroxidation after FLASH-RT compared with CONV-RT is consistent with some recently published in vivo analyses using IF staining of 4-HNE in the lung and gut that suggest less lipid peroxidation in FLASH-RT irradiated samples *(*Vilaplana-Lopera* et al *2025*)*. These results need to be supported by additional quantitative investigations, as measuring lipid peroxidation in tissue is notoriously challenging.

**Figure 7.**
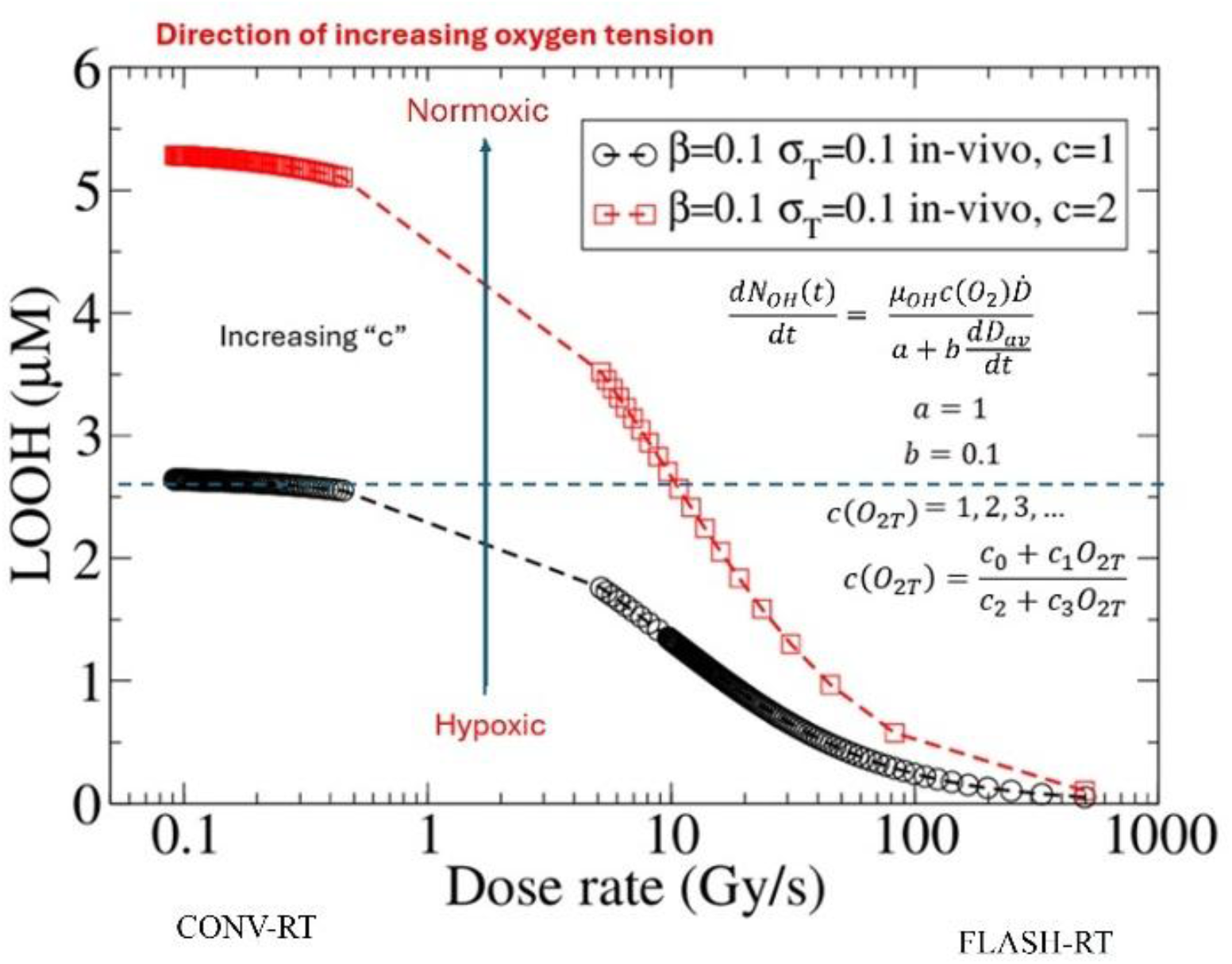
Numerical solutions of the rate equations (Eq. 9-16) for one pulse with a dose of 10 Gy in the brain at different dose rates, showing LOOH production in the brain dependence of dose rate for two different yields of ^•^*OH*.

On the other hand, it is well known that increased lipid peroxidation can damage membranes and tissues (Gaschler and Stockwell 2017). While conventional irradiation–induced cell killing has classically been attributed to DNA damage, increasing evidence indicates that lipid peroxidation could play an important role in normal tissue toxicity *(Pearson et al 2021)*. CONV-RT has shown to increase the protein carbonyl and MDA levels in the brain, reflecting increased protein oxidation and lipid peroxidation *(Gulbahar et al 2009)*. On the other hand, lipid peroxidation in the brain has been proven to have a relevant role in cognitive decline in non-radiation-associated pathologies like Alzheimer’s and vascular dementia *(Ge et al 2026, Zoroddu et al 2026, Bradley-Whitman and Lovell 2015, *Hardas* et al *2013*, *Gustaw-Rothenberg* et al *2010*)*.

The sigmoidal behavior produced by dose rate in lipid peroxidation is also consistent with what has been reported for other biological responses in the long term, such as cognitive assessment after different dose rates, where a sigmoidal behavior was also observed with dose rate, and no significant changes in cognition were observed for dose rates of 10Gy/s and below, and 100Gy/s and beyond *(*Montay-Gruel* et al *2017*)*.

In Figure 7, we can also observe that the differential effect between CONV-RT and FLASH-RT depends on the c parameter of the ^•^*OH* yield as a function of tissue oxygen tension O_2T_. The differential effect of dose rate increases as the ^•^*OH* yield increases. The ^•^*OH* yield depends on the c parameter, which in turn depends on the initial tissue oxygen tension. In that case, the difference between CONV-RT and FLASH-RT on lipid peroxidation would be greater for higher initial tissue oxygen levels and lower as tissue oxygen levels decrease. With this model, we showed that when oxygen perfusion is considered, lipid peroxidation exhibits a sigmoid dose-rate dependence, so some of its biological consequences may also show the same dependence. Additionally, this provides a testable hypothesis as dose-rate-dependent lipid peroxidation should be more easily observed in vivo in normoxic tissues (10 to 40 mmHg) and less readily observed in hypoxic regions (0 to 10 mmHg), such as within clamped normal tissues and tumor tissues. In Figure 7, we can observe that for hypoxic tumors, lipid peroxidation is lower for both CONV-RT and FLASH-RT. Additionally similar amount of LOOH is reached at 0.1 Gy/s for hypoxic tumors and at 10 Gy/s for normal tissue. In that case, this would be consistent with the FLASH effect observed in vivo, where tumors show no dose-rate dependence *(Vozenin et al 2022, 2026)*. This phenomenon needs to be experimentally validated; however, one possibility is that tumors could respond differently to lipid peroxidation and be less dependent on lipid integrity for survival than normal tissue *(Lei et al 2020)*..

Since LOOH production depends on OH yields, which in turn depend not only on oxygen but also on the total dose, we also expect a total dose effect. This is evident in Fig. 8, where, for both FLASH-RT and CONV-RT, higher doses lead to greater LOOH production; thus, even for FLASH-RT, increasing the dose increases lipoperoxidation, reducing protection. This has also been observed in vivo, where higher doses reduced the protection provided by FLASH-RT *(*Montay-Gruel* et al *2020*)*. Additionally, a trend toward saturation is observed at higher doses, and theoretically, there is a minimal and maximal dose value for each scenario at which the dose-rate effect can be abolished.

**Fig 8.**
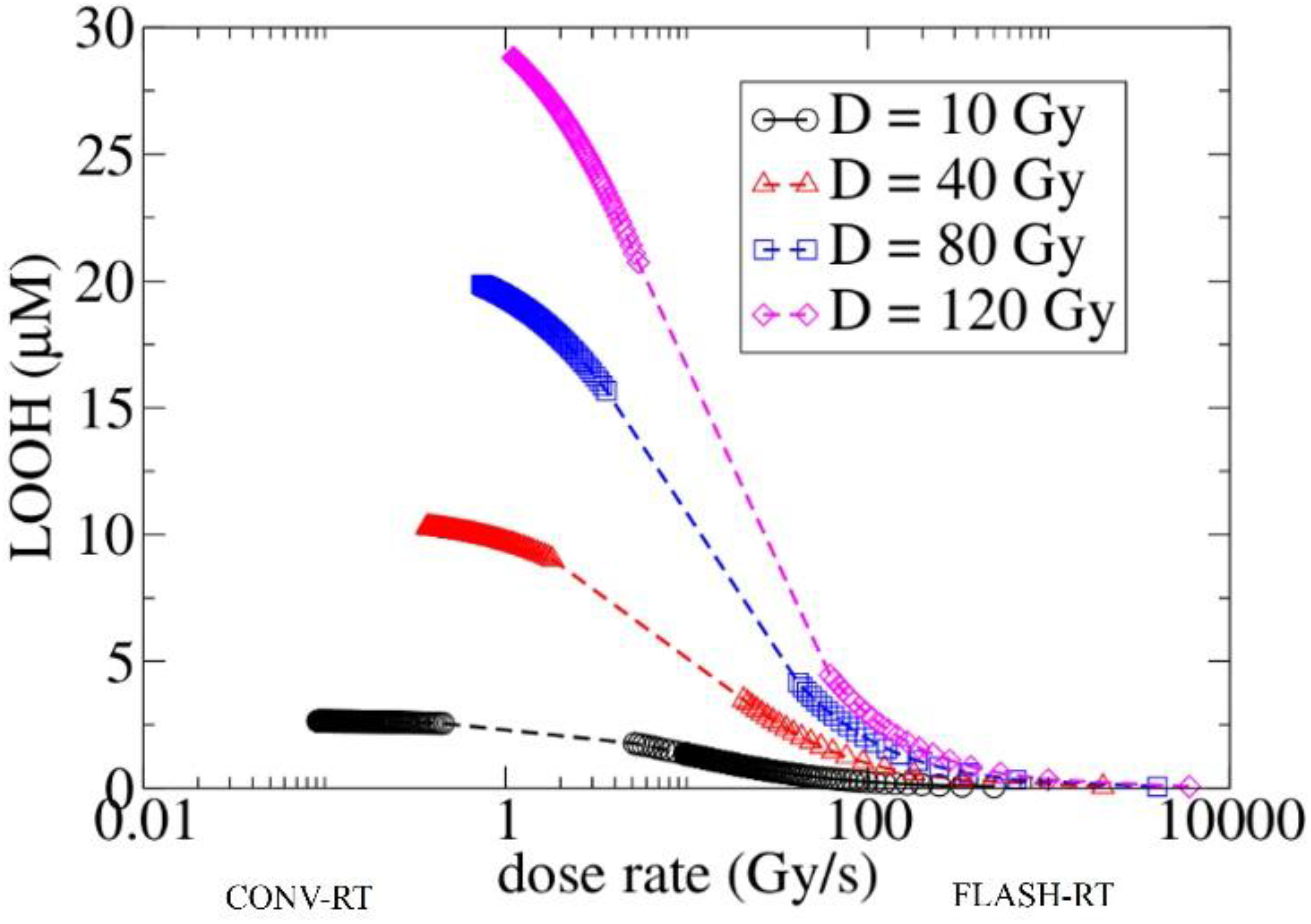
Dose rate dependence of LOOH calculated by the time integration over LOOH for different total dose values. The rest of the parameters are the same as in Figure 7 for c=1. Saturation is observed, as the slope (dashed lines) slows with increasing dose.

### 3.4 How could perfusion and consumption parameters of tissue modify the dose-rate dependence of lipid peroxidation?

We explored how lipid peroxidation varies with the dose rate for different values of oxygen perfusion β, and oxygen consumption *σ*_*T*_, since we also wanted to know what would happen when oxygen reperfusion in the system is modified for tissues other than the mouse brain or for different experimental models. We first calculated lipid peroxidation in the absence of perfusion (β = 0; Fig. 9). When perfusion is stopped, the maximum difference between FLASH-RT and CONV-RT decreases because lower perfusion diminishes the dose-rate effect. Without oxygen perfusion, we can also see that at a very slow dose rate, oxygen consumption and irradiation could permanently deplete oxygen, resulting in less lipid peroxidation. The lack of oxygen reperfusion and ongoing oxygen consumption can mimic some in vivo clamping experiments, limiting blood circulation, and even mimic regions within tumors with minimal oxygen perfusion. In subcutaneous tumors, clamping experiments are known to induce tumor resistance to CONV-RT, however recently we showed that acute hypoxia does not modify tumor response to FLASH-RT (Leavitt *et al* 2024). These results are consistent with Figure 9, where reducing perfusion has minimal impact on FLASH-RT LOOH production, while there is a significant effect observed for CONV-RT. Additionally, the simulation (Fig. 9) shows that when perfusion is absent (b=0, red dots),some extremely low and extremely high dose rates can yield the same LOOH production values. So when perfusion is absent the more extreme dose-rates we compare the smallest dose-rate effect, whereas differences may exist when comparing intermediate CONV-RT values with FLASH-RT. For the dose rates reported in Leavitt et al. (marked in circles in Fig. 9) and no perfusion (red dots), we observe that FLASH-RT produces slightly higher lipid peroxidation than CONV-RT. This is consistent with a slightly increased doubling tumor time observed after FLASH-RT for clamped tumors (no perfusion) (Leavitt *et al* 2024).

**Fig 9.**
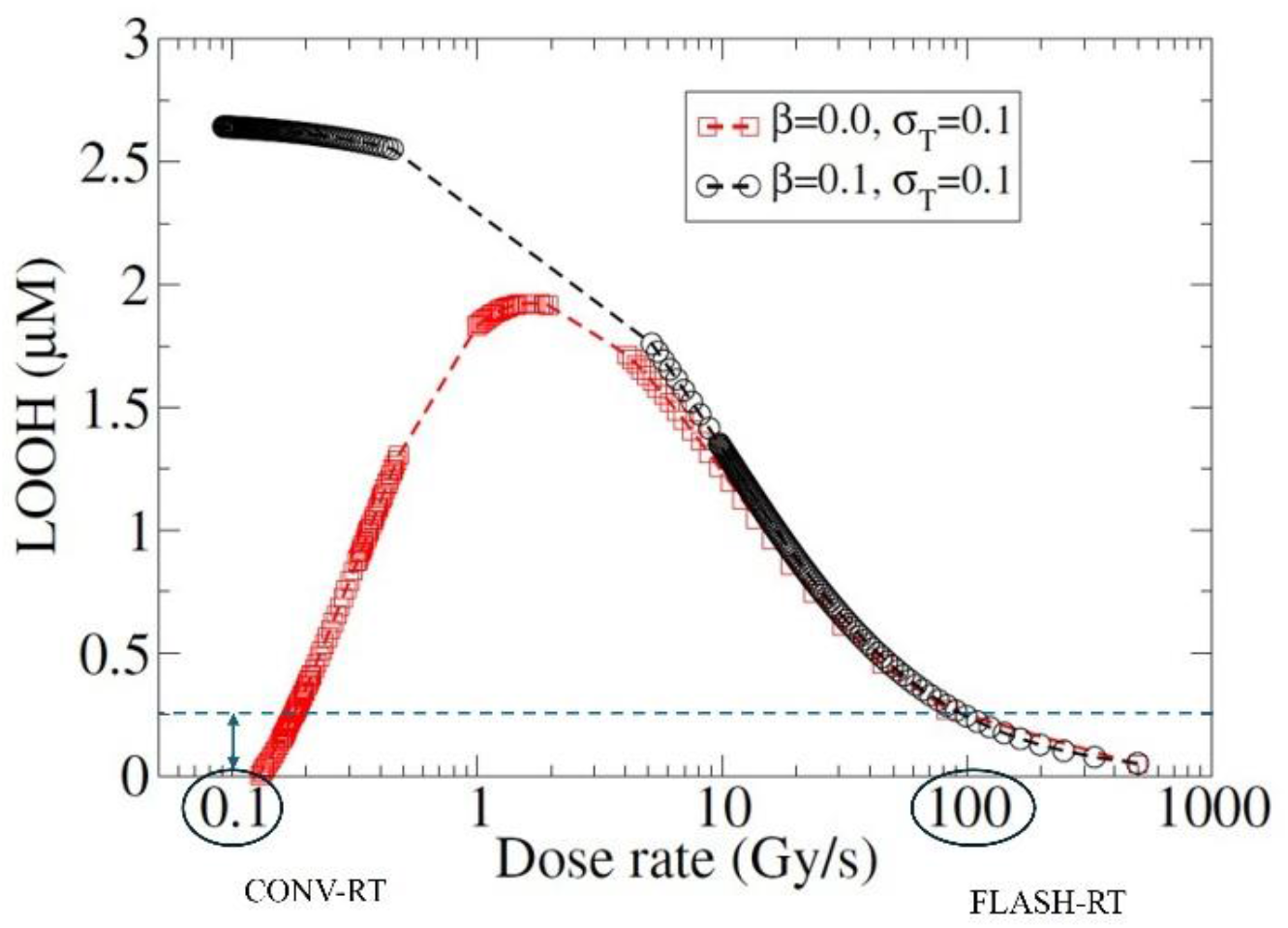
Dose rate dependence of LOOH calculated by the time integration over LOO*H* for different perfusion β values. Circles mark the dose rates used for clamping experiments in Leavitt et. al. (Leavitt *et al* 2024)

Although this model was not intended for that, continuous consumption but no perfusion, could also resemble oxygen dynamics observed in in vitro experiments, where oxygen diffusion in cell cultures under ambient conditions (21% O2) can slow by two or more orders of magnitude compared to in vivo, since diffusion cannot compensate for consumption beyond the surface (Al-Ani *et al* 2018a). Our results indicate that when perfusion is reduced while consumption is maintained, the difference between CONV-RT and FLASH-RT can be reduced. CDRT (Nomura *et al* 2024, Adrian *et al* 2020).

Furthermore, when considering a scenario with no perfusion and no physiological oxygen consumption (β = *σ*_*T*_ ≈ 0), a dose rate dependence in lipid peroxidation becomes evident again, showing a distinct difference between CONV-RT and FLASH-RT as illustrated in Figure 10 (black curve). This pattern resembles the behavior observed in liposomes in water, where perfusion is low, and no oxygen consumption occurs. (Froidevaux et al., 2023).

**Fig 10.**
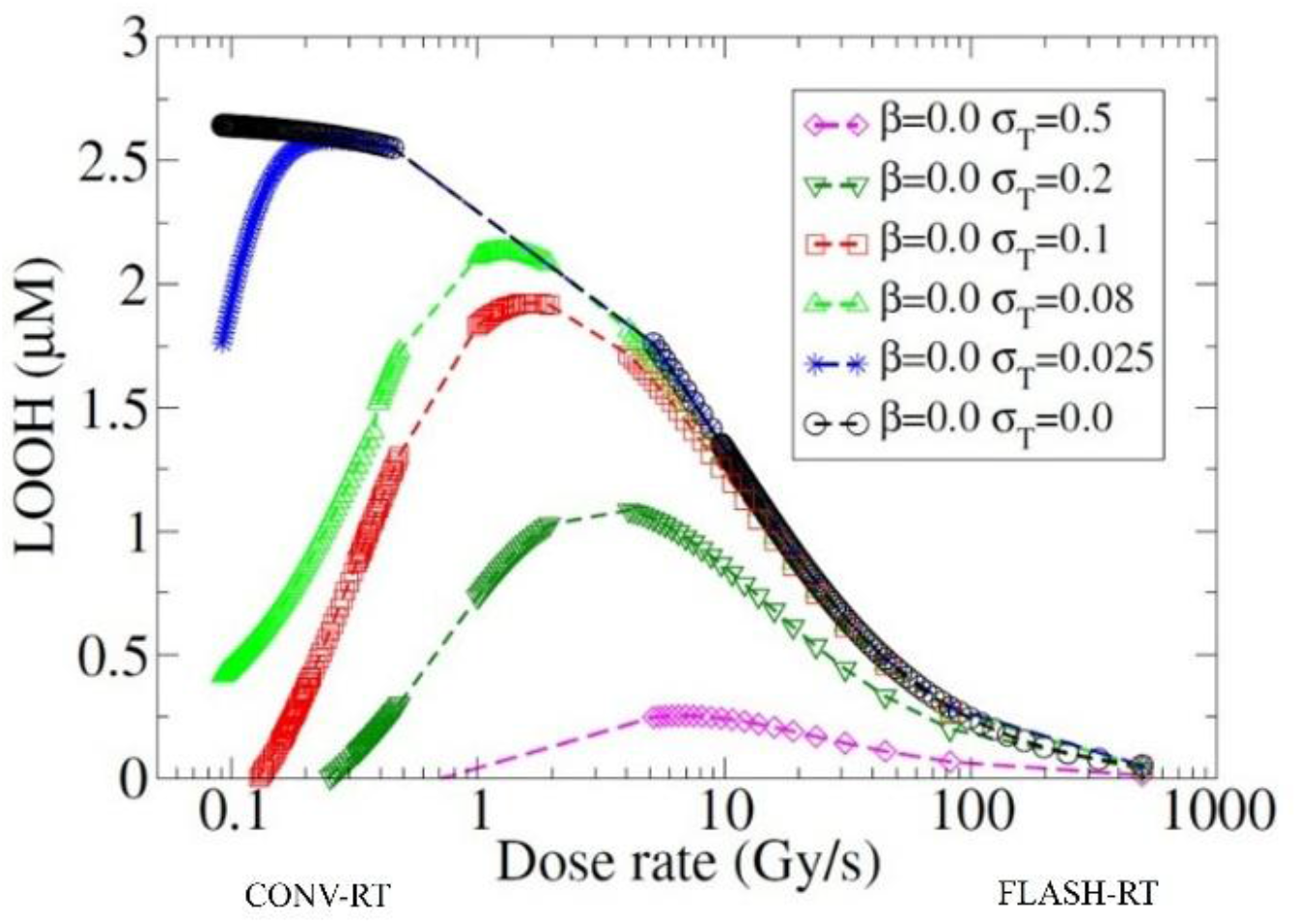
Dose rate dependence of LOOH calculated by the time integration over LOO*H* for no oxygen perfusion β= 0 and different oxygen consumption σ_T_ values with initial PO_2_ 40mmHg.

Interestingly, also in Figure 10. We can observe that for the case where there is no perfusion, but consumption is still very high, the dose rate effect in lipid peroxidation gets more flattened. If we assume that oxygen perfusion and consumption parameters vary between tumor and normal tissues, we can propose that the dynamics of oxygen perfusion can create a dose-rate-dependent effect on oxygen-dependent reactions, such as lipid peroxidation, depending on these parameters.

Finally, in Fig. 11, we again observe that, for the same case without reperfusion but with physiological consumption values (resembling clamping experiments or tumor regions), the higher the initial oxygenation (*O*_2*T*_|_*T*=0_), the greater the difference between FLASH-RT and CONV-RT. In contrast, in tissues with very low initial oxygen tension, lipid peroxidation is absent, and there is no difference between dose-rate modalities. We also observe a saturation effect as initial oxygen levels increase. This graph may not fully represent in vitro settings, since when the initial oxygenation values are modified, generated gradients could immediately alter the reperfusion parameters, depending on the experimental conditions (Al-Ani *et al* 2018b).

**Fig. 11.**
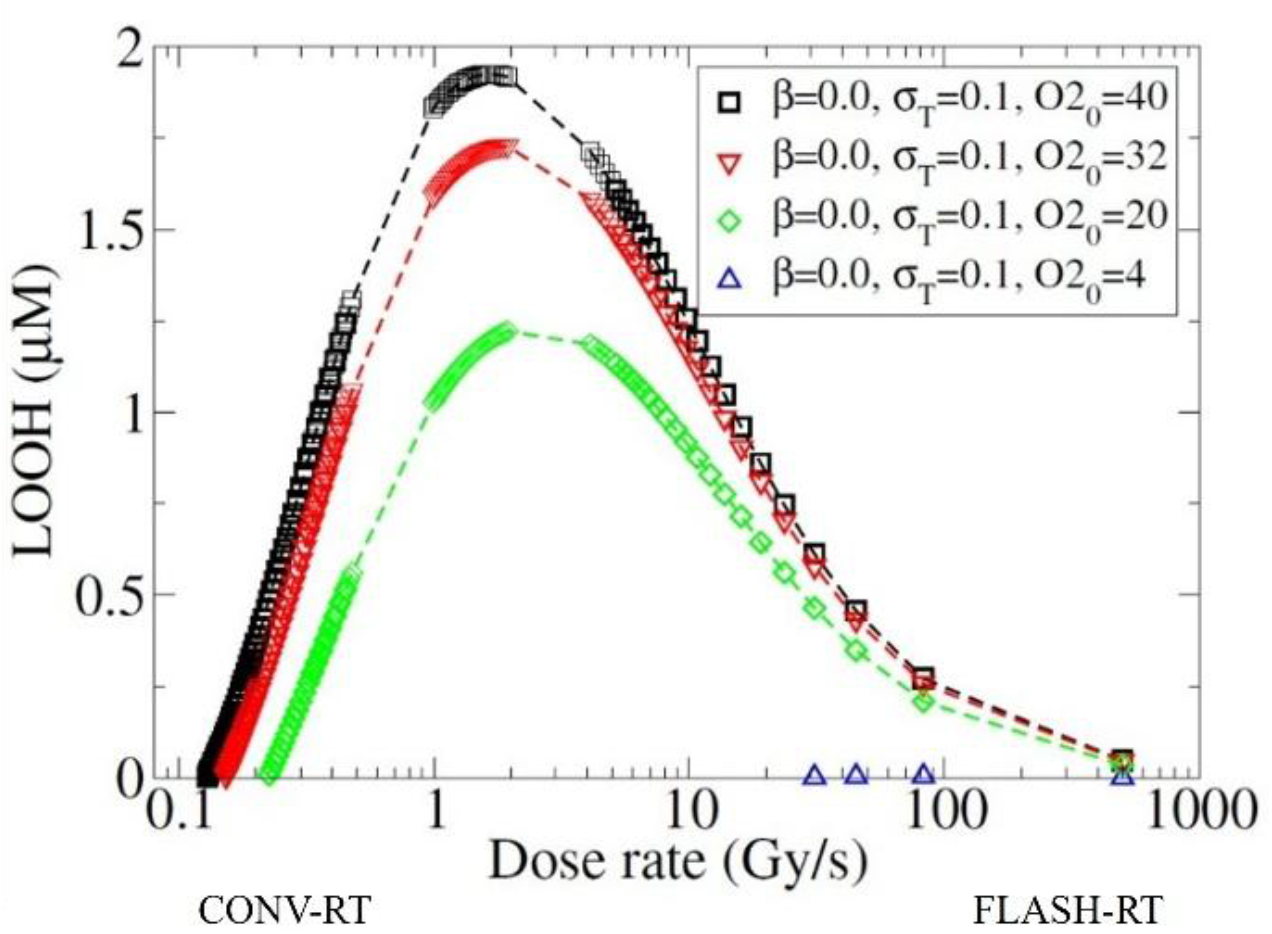
Dose rate dependence of LOOH calculated by the time integration over LOO*H* for no oxygen perfusion, β= 0, and consumption σT=0.1 for different oxygenation values.

### 3.5 The Mexican Subway hypothesis

Considering the role of oxygen perfusion in tissues, FLASH-RT can be compared to rush hour on a crowded subway, where many people arrive simultaneously and all board the same train. All physical interactions involving a defined absorbed dose, like ionization, aqueous electrons, and reactive species produced occur and arrive to the tissue almost simultaneously —equivalent to a defined number of people entering the same train at the same time — reacting subsequently with local oxygen existing in the tissue within that brief moment —comparable to all people entering in the train and breathing from the existing oxygen inside the train—. In contrast, conventional dose rates resemble people arriving gradually over an extended period, each taking different trains. This staggered arrival allows for oxygen to be replenished by each train, —comparable to blood oxygen transport and oxygen replenishing occurring in tissue—. As a result, this initial interaction of people with oxygen upon entering the train is significantly higher in conventional dose rates than during rush hour or, in our case, during FLASH-RT. Therefore, the time frame of ionizing radiation arrival and interaction with tissue may determine its interaction with oxygen perfusion (train arrivals) (Fig. 12B). These temporal oxygen dynamics may be relevant to subsequent biological responses in tissues, as they can influence oxygen-dependent reactions, such as lipid peroxidation, which initiate within a specific time frame. So the faster the arrival, the less interaction with oxygen perfusion. Therefore, oxygen dynamics in the tissue could generate a dose-rate-dependent response in the long term.

**Fig 12.**
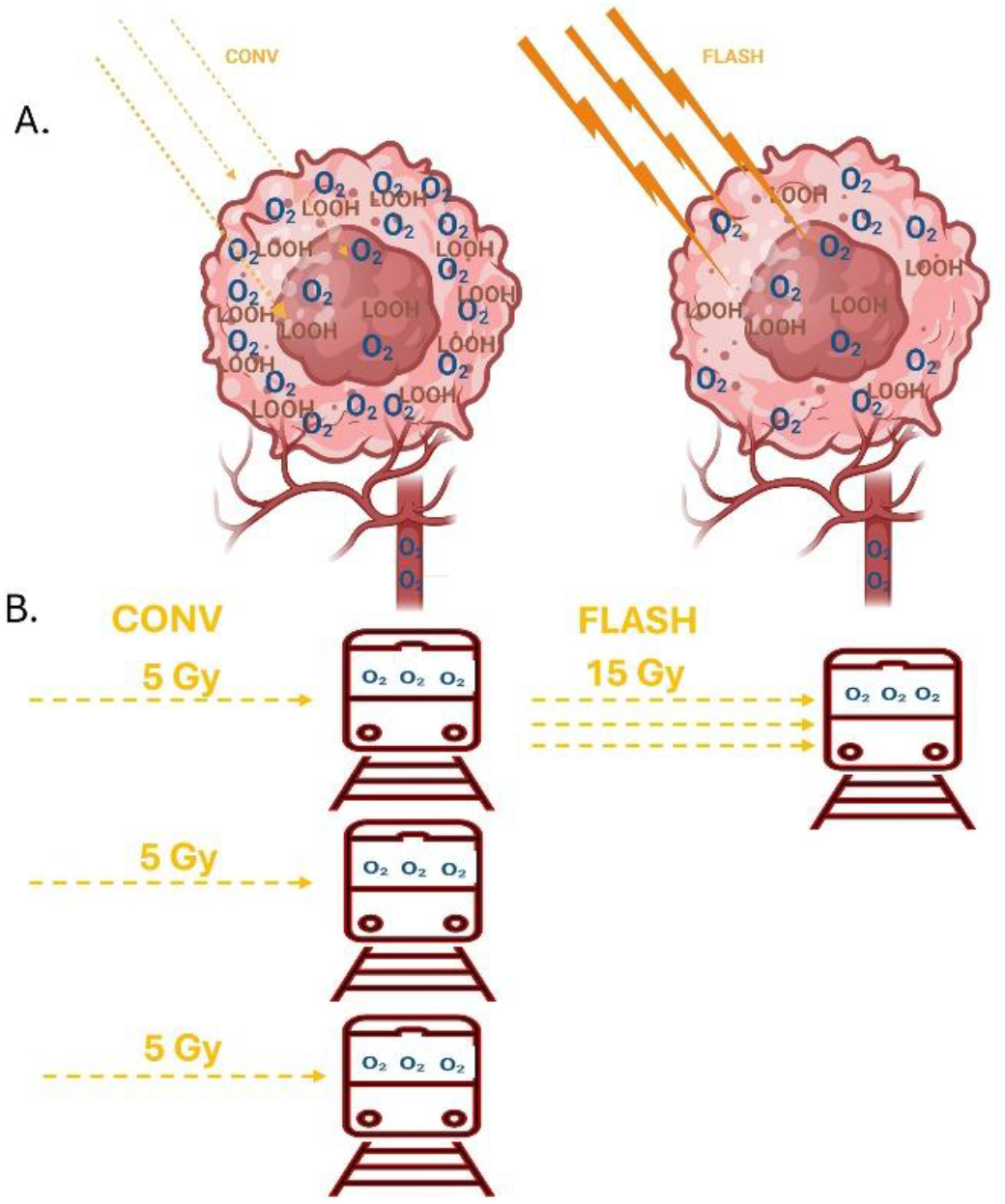
Scheme showing how differences in tissue oxygenation dynamics can influence dose rate dependence in lipid peroxidation. A. The scheme shows that differences in oxygenation between tumor tissues at the center and well-perfused normal tissues would lead to differences in lipid peroxidation, and how the dose rate would affect only the well-perfused tissues. B. The scheme shows subway trains representing oxygen delivery via perfusion and how the same radiation dose interacts with more oxygen at CONV-RT than at FLASH-RT (Created in https://BioRender.com).

## Conclusions

This compartmental model has proven effective in demonstrating how tissue oxygen perfusion may contribute to dose-rate-dependent effects. It is important to highlight the fact that the influence of dose rate on lipid peroxidation is closely related to the initial oxygenation levels. Our calculations indicate that as oxygenation decreases, the dependence on dose rate may also decrease. Moreover, we have identified that the relationship between dose rate and lipid peroxidation can be affected by oxygen perfusion and consumption parameters across different tissues. In particular, for tissues characterized by minimal oxygen perfusion, high oxygen consumption, and high metabolic demands—such as tumors or epidermis—we anticipate observing a reduced effect of dose rate on lipid peroxidation. The in vivo oxygen perfusion dynamics may therefore impact the levels of oxygen-dependent radiochemical reactions and downstream biological responses. These first simulations were performed under the assumption of a continuous beam. Future simulations will investigate the in vivo interaction between the temporal dynamics of oxygen perfusion and the temporal structure of pulsed electron beams. Ultimately, we propose that temporal oxygen dynamics in tissues could be a relevant mechanism contributing to the biological effects of FLASH.

## Acknowledgements

Funding was provided by Swiss National Science Foundation grant Spirit IZSTZ0_198747/1 (to MCV and PBZ supporting JFC); American Cancer Society Diversity in Cancer Research Institutional Development Grant (DICRIDG-21-074-01-DICRIDG) at Howard University (to RA); MAGIC-FNS CRSII5_186369 (to MCV and supporting VG); and CONACYT for supporting PBZ’s sabbatical in Switzerland. Finally, we would like to thank the core facilities and staff at Lausanne CHUV, as well as their veterinary staff, for their valuable support.

